# ATM functions as a rheostat of metabolic stress in small-cell lung cancer

**DOI:** 10.64898/2026.03.13.711672

**Authors:** Debdatta Halder, Utsav Sen, Vrinda Jethalia, Subhamoy Chakraborty, Andrew Elliott, Kedwin Ventura, Ari Vanderwalde, Balazs Halmos, Hossein Borghaei, Tin Htwe Thin, Alan Soto, Mirela Berisa, Rachel Brody, Deniz Demircioglu, Dan Hasson, Triparna Sen

## Abstract

ATM is best known as a guardian of genomic stability, yet its contributions to oncogenic signaling in aggressive malignancies like small-cell lung cancer (SCLC) remain poorly understood. Despite ATM being an established clinical vulnerability in SCLC, its influence on dysregulated tumorigenic circuits remains unclear. We demonstrate that inhibition of ATM disrupts the AKT-mTORC1-4EBP1 signaling axis, leading to attenuation of the master regulator of stress, ATF4. ATF4 and MYC appear to co-regulate one another in a feedback loop critical for redox homeostasis. ATM inhibition perturbs both the expression and function of MYC and ATF4, leading to increased intracellular reactive oxygen species, impaired glutathione recycling, and ferroptotic cell death, thereby exposing a crucial dependency of SCLC on stress-adaptive signaling. We uncover previously unrecognized metabolic vulnerability in SCLC, nominating ATM as a regulator of adaptive stress, expanding its role beyond canonical DNA damage repair (DDR) and highlighting therapeutically exploitable opportunities in aggressive tumors.

**Statement of Significance:** The metabolic landscape of SCLC remains poorly characterized, particularly its interaction with dysregulated signaling networks, limiting the development of effective strategies to overcome therapeutic resistance. Our work reveals an expanded role for ATM beyond DNA repair, positioning it as a key regulator of metabolic rewiring and highlighting new therapeutic opportunities for SCLC.

## Introduction

Small-cell lung cancer (SCLC) is the most aggressive form of lung malignancy, accounting for approximately 15% of all lung cancer cases and around 30,000 new diagnoses annually in the United States.^1,2^ Despite its initial sensitivity to chemotherapy, generally the disease quickly relapses, and the median overall survival (OS) remains dismal at less than a year.^2^ The 5-year survival rate is approximately 7%, underscoring the urgent need for novel therapeutic strategies.^1^ Standard first-line treatment has long relied on platinum-etoposide chemotherapy, with or without immune checkpoint inhibitors, but durable responses are rare and resistance emerges rapidly. Although recent efforts have introduced surface-targeted agents (DLL3-targeted therapies), enhancing prognosis in the second-line treatment setting^3^, these therapies are not curative and have demonstrated severe toxicities,^4^ limiting their clinical benefit. Therefore, it is critical to uncover targetable vulnerabilities for SCLC. A particularly understudied area in SCLC biology is its metabolic profile. Current knowledge of maladaptive signaling in SCLC and its crosstalk with metabolic pathways is insufficient. Exploitation of these functional circuits can help target drug resistance efficiently.

DNA damage response (DDR) has been demonstrated to be a potent therapeutic target in SCLC due to its inherent genomic instability, high replication stress, and near-universal loss of *TP53* and *RB1*.^6^ Consequently, inhibitors of key DDR proteins, such as PARP, CHK1, and ATR, have shown preclinical efficacy and are under clinical evaluation in biomarker-selected and unselected subsets of patients.^7–10^ Central to this process is the ATM kinase, a key phosphatidylinositol 3-kinase-related kinase (PIKK) family member that activates downstream repair, checkpoint, and apoptotic pathways in response to DNA double-strand breaks (DSBs).

Although ATM traditionally has been recognized to function principally as a guardian of genome integrity, recent studies portray their additional critical noncanonical roles in maintaining cellular homeostasis. In particular, ATM helps regulate oxidative stress responses, influencing mitochondrial function, metabolic pathways, and even gene expression programs beyond its classic DDR function.^11–16^ ATM is activated under hypoxic conditions where it phosphorylates and stabilizes HIF-1α, thereby enhancing cancer cell survival in low-oxygen environments.^17^ In breast cancer models, oxidative activation of ATM in cancer-associated fibroblasts drives a glycolytic switch that fuels tumor cell invasion, highlighting a metabolic coupling mechanism.^18^ Additionally, in response to oxidative stress, ATM phosphorylates p53 at Ser15, which can either promote apoptosis or induce antioxidant programs via sestrins, GPX1, and MnSOD.^19^ ATM is essential for full activation of the AKT pathway in response to DNA damage, including AKT phosphorylation at Ser473^20,21^ and GLUT4 translocation to the cell surface, which enhances glucose uptake.^22^ This signaling cascade activates the pentose phosphate pathway, increasing NADPH production and GSH recycling through G6PD upregulation, both critical for redox control and anabolic metabolism.^11^ Metastatic breast cancer cells with elevated AKT activity are particularly dependent on this axis, and ATM inhibition triggers apoptosis.^23^ Beyond metabolic regulation, ATM can promote epithelial-to-mesenchymal transition (EMT), cancer cell invasion and dissemination^24^ by activating NF-κB signaling through nuclear export and cytoplasmic relocalization in breast cancer.^25^ Collectively, these observations suggest that in cancers with defective cell cycle checkpoint responses, elevated ATM activity may serve as a central hub for metabolic adaptation, redox balance, and pro-metastatic signaling.^24^ This multifaceted role has been described in other cancers,^26^ but it is largely unknown how ATM can modulate oncogenic signaling and metabolic rewiring in *TP53* and *RB1*-deficient aggressive tumors like SCLC that harbor chronic replication stress, relying on constitutive stress-response pathways for survival.^27^

We demonstrate that ATM can regulate oncogenic signaling via the AKT-mTORC1-ATF4 axis and preserve redox homeostasis of the cell via a supportive amino acid pool, harnessing a MYC-mediated transcriptional rewiring and preventing cell death. In addition, we demonstrate that ATM inhibition causes ferroptosis-mediated cell death and antitumor response in SCLC. Our findings extend the function of ATM beyond genome maintenance, highlighting it as a central gatekeeper of metabolic adaptation and protein homeostasis in SCLC, uncovering a novel biology and therapeutic strategy for refractory SCLC.

## Results

### High ATM expression predicts poor prognosis and resistance to chemotherapy and other DDR agents in SCLC

To assess whether ATM has a functional role in SCLC biology and patient outcomes, we conducted a comprehensive analysis of transcriptomic data from 174,226 clinical specimens spanning 23 cancer types. Notably, SCLC demonstrated the highest *ATM* gene expression among all tumor types analyzed (*n* = 944 SCLC samples; **Fig. 1A**); within lung cancer, *ATM* expression was highest in SCLC compared to non-small-cell lung cancer (NSCLC), large-cell lung cancer, lung carcinoid, and atypical carcinoid (**Fig. 1B**). Details are listed in **Supplementary Table S1**.

**Figure 1.**
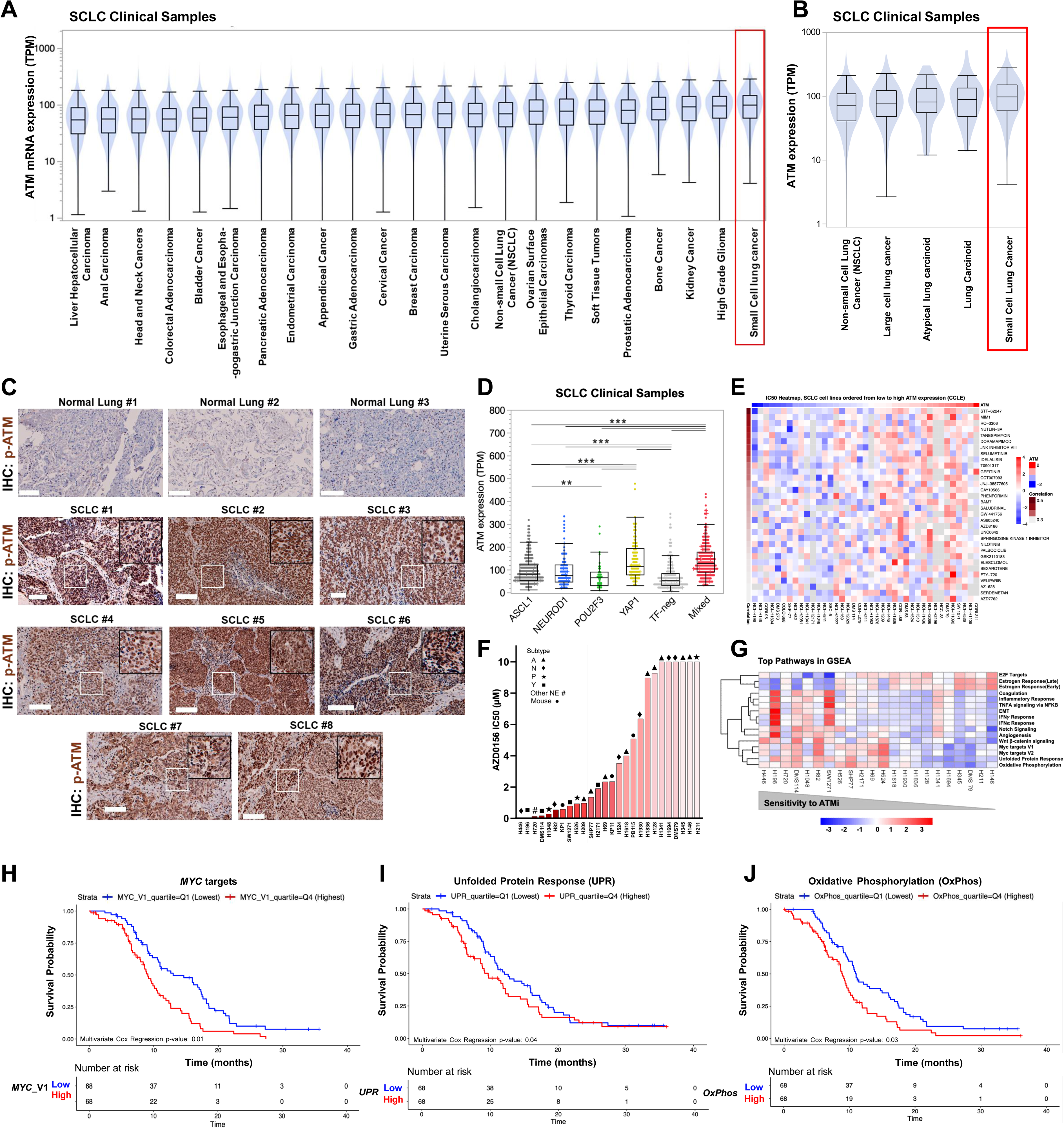
ATM is highly expressed in SCLC patients and mechanistically concordant with worse prognosis. **(A)** Comparative mRNA expression of ATM in different cancer tissues collected from 174,226 real-world patient tumor samples subjected to RNA-seq. **(B)** Boxplots showing comparative mRNA expression of ATM in 5 major types of lung cancer, using the same dataset as in **(A)**. SCLC is highlighted in a red box (*n* = 944) in both **(A)** and **(B). (C)** IHC of real-world patient tissue with phospho-ATM (Ser1981), the active form of ATM, across 3 normal lung and 8 SCLC tissue samples. Scalebar = 100 µm. **(D)** SCLC subtype-wise comparison of mRNA expression of ATM in real-world patient data (*n* = 944 SCLC clinical samples). Subtypes are driven by relatively high expression of TFs *ASCL1*, *NEUROD1*, *POU2F3*, and *YAP1*, 1 or more of them (mixed), or TF-negative (low expression of all 4 TFs). **(E)** Heatmap showing *Z*-score-normalized IC_50_ values of 39 SCLC cell lines (heatmap columns) to various anticancer drugs (CCLE database). Cell lines are ordered from low-to-high ATM expression (as seen in the column annotation bar above). Spearman correlation coefficients between ATM expression and drug sensitivity are displayed as a row annotation bar. **(F)** IC_50_ values of 26 SCLC cell lines representing all major subtypes of SCLC treated with ATMi AZD0156. **(G)** Gene set enrichment analysis (GSEA) pathway analysis showing top pathways of sensitivity in response to AZD0156. Heatmap shows *Z*-scored average expression of leading-edge genes of the pathways. Cell lines are ordered based on high-to-low sensitivity to AZD0156 (low-to-high IC_50_). **(H-J)** Kaplan-Meier survival analysis of patients stratified by top GSEA signatures associated with high sensitivity to AZD0156 in **(G).** MYC target gene expression (Myc_Targets_V1) **(H),** UPR **(I),** oxidative phosphorylation **(J)** represented in the IMpower133 dataset; only patients in the top quartile (Q4; red) and bottom quartile (Q1; blue) of signature expression are displayed for OS. High signature expression was significantly associated with poorer survival outcomes as statistically tested on the entire dataset of 271 patients (multivariate Cox regression *P* = 0.01, *P* = 0.04, and *P* = 0.03 for **H, I,** and **J,** respectively). The number of patients at risk at each time point is shown below the graphs.

We next validated ATM protein expression levels using immunohistochemistry (IHC) in SCLC clinical samples. Compared to normal lung tissue, SCLC tumors showed markedly elevated levels of both total ATM **(Supplementary Fig. S1A**) and the active phosphorylated form (pATM Ser1981) (**Fig. 1C**), indicating both increased abundance and activation of ATM in SCLC. Importantly, *ATM* gene expression and protein levels were only weakly correlated (**Supplementary Fig. S1B**), suggesting the need for multimodal profiling when assessing ATM status in clinical samples. *ATM* gene expression was comparable across primary and metastatic SCLC tumors, suggesting consistent *ATM* expression throughout disease progression (**Supplementary Fig. S1C, Supplementary table S1**).

We further examined the clinical relevance of *ATM* expression in SCLC. In a patient cohort treated with cisplatin-based chemotherapy (*n* = 126), patients with high *ATM* expression had worse OS than those with low *ATM* expression (HR = 1.715; **Supplementary Fig. S1D**). These findings highlight *ATM* as a potential biomarker of poor prognosis and chemotherapy resistance in SCLC.

To further contextualize *ATM* expression within the molecular heterogeneity of SCLC, we stratified tumors by transcription factor (TF)-based subtypes (*ASCL1*, *NEUROD1*, *POU2F3*, *YAP1*, TF-negative, and mixed). *ATM* expression was highest in the YAP1-positive and mixed-TF subtypes (**Fig. 1D, Supplementary Table S1**).

Our analysis reveals that *ATM* is markedly upregulated and activated in SCLC compared to other cancer types and lung cancer subtypes. High *ATM* expression is associated with poor clinical outcomes, particularly reduced OS following chemotherapy, establishing ATM as a potential prognostic biomarker in SCLC.

### High ATM expression predicts drug resistance, and SCLC models are sensitive to ATM inhibition

Given the canonical role of *ATM* in DNA repair, we next examined the association of *ATM* expression with other DDR genes across SCLC samples. Among the 944 clinical specimens analyzed, *ATM* expression was strongly correlated with multiple DDR-related genes, including *PRKDC* (Spearman ρ = 0.82), *ORC6* (ρ = 0.78), and *MTF2* (ρ = 0.76) (**Supplementary Fig. S1E**). This reinforces ATM’s role as a central player in the DDR networks in SCLC. Next, to investigate whether ATM contributes to resistance beyond cisplatin, we analyzed drug response data from 39 SCLC cell lines in the Cancer Cell Line Encyclopedia (CCLE). High ATM expression correlated with resistance (higher half-maximal inhibitory concentration [IC_50_]) to 32 drugs, including AZD7762, a dual Chk1 and Chk2 inhibitor (**Fig. 1E**). This suggests that ATM may broadly be associated with resistance to DNA-damaging and checkpoint-targeting agents.

We then explored the therapeutic potential of targeting ATM directly in SCLC. Using 2 independent ATM inhibitors (ATMis) (AZD0156 and KU55933), we tested a panel of 26 SCLC cell lines representing all major molecular subtypes. AZD0156 treatment inhibited growth in 20 of 26 cell lines, with 12 cell lines exhibiting IC_50_ values < 2 μM (**Fig. 1F**). KU55933, though less effective than AZD0156, confirmed the sensitivity of SCLC lines to ATM inhibition, further supporting ATM as a druggable vulnerability (**Supplementary Fig. S1F**).

Mechanistic studies revealed that ATMi-induced G2/M cell cycle arrest in H446, H196, and DMS114 cell lines (**Supplementary Fig. S1G**). Furthermore, pharmacological inhibition of ATM caused apoptosis that was partially rescued by the caspase inhibitor ZVAD-FMK (**Supplementary Fig. S1H**). These results indicate that ATM is essential for cell cycle progression and survival in SCLC cells, consistent with previous reports emphasizing the dependence of SCLC on DDR at the G2/M cell cycle checkpoint.^27^

Given the well-known role of PARP1 in SCLC and ATMi resistance, we next investigated whether combination therapy targeting both ATM and PARP1 could overcome resistance. We treated 3 SCLC cell lines (H196, H446, and DMS79) with AZD0156 (ATMi) and AZD5305 (a selective PARP1 inhibitor currently in clinical trials). Combination therapy yielded greater cytotoxicity compared to either agent alone in all 3 models (**Supplementary Fig. S1I-S1K**). In H446 cells, the combination IC_50_ was 2.6 nM versus 150 nM and 930 nM for AZD0156 and AZD5305 monotherapies, respectively (**Supplementary Fig. S1J**). These findings demonstrate a synergistic interaction between ATM and PARP1 inhibition in SCLC models (**Supplementary Fig. S1I-S1K**).

Most SCLC cell lines, including diverse molecular subtypes, are sensitive to pharmacologic ATM inhibition, and combined targeting of ATM and PARP1 produces synergistic cytotoxicity, supporting a combinatorial treatment strategy.

### ATM expression correlates with DDR activity and predicts vulnerabilities in specific molecular contexts

To identify potential biomarkers of response to ATM inhibition, SCLC cell lines were ranked by their sensitivity to AZD0156 (IC_50_ values), and pathway-level enrichment analysis was performed on their baseline transcriptomic profiles (**Fig. 1G**). SCLC lines with high sensitivity (IC_50_ < 2 μM) displayed elevated expression of *MYC* target genes, oxidative phosphorylation, and unfolded protein response (UPR) pathways. As cell lines with high ATMi sensitivity showed high expression of these pathways, the data suggest that these cellular programs may be the major downstream mediators of ATM dependency in SCLC.

To determine whether these pathways have prognostic significance in patients with SCLC, we analyzed the IMpower133 cohort of 271 SCLC cases.^28^ IMpower133 provides high-quality, uniformly treated survival data, ideal for prognostic analysis. High expression of *MYC* targets, oxidative phosphorylation, and UPR genes each predicted significantly decreased OS (**Fig. 1H– J**). Conversely, resistant cell lines (IC_50_ > 10 μM) were enriched for *E2F* target gene expression, pointing to distinct transcriptional programs that may drive differential drug responses (Fig. 1G).

Hence, *ATM* expression strongly correlates with DNA repair gene networks, and sensitivity to ATM inhibition in SCLC is associated with a subset of tumors with high expression of *MYC* targets, oxidative phosphorylation, and UPR pathways, highlighting possible noncanonical roles of ATM in SCLC pathobiology.

### Genome-wide CRISPR/Cas9 knockout screen demonstrates that ATM inhibition disrupts AKT-mTORC1 signaling and suppresses the unfolded protein response in SCLC

To identify downstream mechanisms by which ATM inhibition decreases SCLC viability, we performed genome-wide CRISPR/Cas9 knockout screens in the highly sensitive H196 SCLC cell line. For these screens, H196 cells were first infected with the genome-wide TKOV3 sgRNA library (Toronto Knockout CRISPR Library-Version 3) containing 70948 sgRNAs across 18,053 protein-coding genes at a multiplicity of infection (MOI) of 0.3 (see ***Methods***). On day 5 (T0), all cells were pooled and split into 2 treatment arms: 1) control and 2) ATMi, AZD0156 (20% inhibitory concentration [IC_20_]). After 1 doubling (T1) and 3 doublings (T2), cells were harvested for sequencing (**Supplementary Fig. S2A**). As expected for a positive selection screen, sgRNA enrichment/depletion comparing biological replicates showed that the replicates were modestly correlated with robust enrichment of some sgRNAs in 3 replicates for each condition. After treatment with AZD0156, the PI3K/AKT pathway was found to be a top enriched pathway at the last doubling (T2), suggesting that the PI3K/AKT pathway is necessary for AZD0156 cytotoxicity (**Supplementary Fig. S2A, Fig. 2A**).

**Figure 2.**
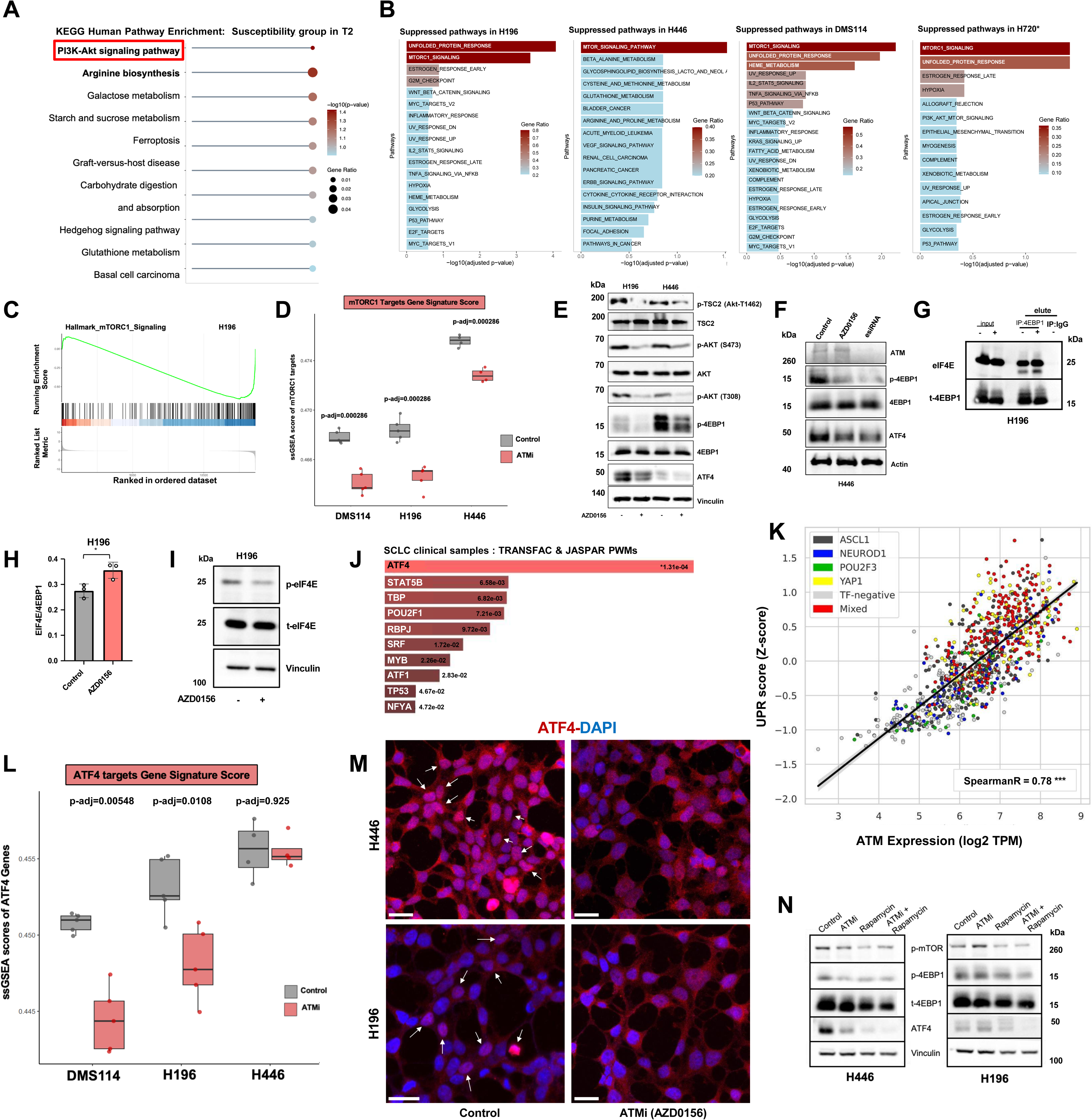
ATM inhibition downregulates the integrated stress response through inhibition of AKT and mTORC1. **(A)** Functional enrichment analysis of top-ranked genes identified from the CRISPR screen. Genes selected from the upper end (n=42 genes) of the sigmoid-ranked essentiality curve were analyzed for pathway enrichment. Lollipop plot displays the top depleted biological processes at T2, with statistical significance indicated by -log₁₀ (nominal *P*-value) (color scale) and gene ratio (dot size). Enriched terms reflect biological processes preferentially required under ATM inhibition (AZD0156). **(B)** RNA-seq over representation analyses from MSigDB Hallmark gene sets and KEGG databases of differentially expressed genes following AZD0156 treatment in 4 SCLC cell lines. **(C)** GSEA plot showing the top negatively enriched pathway across 4 SCLC models treated with AZD0156 compared to control, represented in H196. The green line indicates the enrichment score (ES), and vertical black bars represent the position of mTORC1 signature genes in the ranked gene list. **(D)** Boxplots showing mTORC1 target gene signature scores in SCLC cells treated with ATMi AZD0156 (red) compared to DMSO controls (gray) across 3 SCLC cell lines: DMS114, H196, and H446. *P*-values were calculated using a 2-sample *t*-test and were adjusted using the false discovery rate (FDR) method (*P*-adj = 0.000286 for all comparisons). **(E)** Immunoblot assay of H446 and H196 +/- AZD0156, probed for AKT-mTORC1 pathway markers (p-T1462-TSC2, TSC2, p-T308-AKT, AKT, p-4EBP1, 4EBP1) and UPR/ISR marker ATF4. **(F)** Immunoblot of H446 cell lysates treated with either AZD0156 (ATMi) or ATM siRNA pool (esiRNA) and probed for the major markers p-4EBP1 and ATF4. ATM was visualized to evaluate the efficacy of the siRNA pool. Actin or vinculin was visualized as a loading control **(E-F). (G-H)** Immunoblot of eIF4E and total-4EBP1 after H196 cells were treated with or without AZD0156 (72 h). Whole-cell lysates were immunoprecipitated with 4EBP1 and eluted with low-pH glycine buffer (elute). Basal expressions were visualized in whole lysate (input). Boxplot shows the quantified ratio of eIF4E bound to 4EBP1 in the elute **(H)**. *P*-value was calculated by Student’s *t*-test. **(I)** Immunoblot of H446 and H196 +/- ATMi probed for p-eIF4E, eIF4E, and vinculin. Vinculin was used as a loading control. **(J)** TF binding motif enrichment (JASPAR and TRANFAC PWMs analysis) from the top 50 genes positively correlated to ATM expression in real-world SCLC patient samples (*n* = 944). **(K)** Scatterplot representation of Spearman correlation between ATM mRNA expression and UPR score across real-world patient SCLC tumor samples from different transcriptional subtypes. Each dot represents an individual tumor, color-coded by subtype: ASCL1 (blue), NEUROD1 (black), POU2F3 (green), YAP1 (yellow), TF-negative (gray), and mixed (red). The black line indicates the best-fit regression line. Spearman correlation coefficient R = 0.78 (\*\*\**P* < 0.001), *n* = 944 SCLC patients. **(L)** Boxplots showing ATF4 target gene signature scores in SCLC cells treated with ATMi AZD0156 (red) compared to DMSO controls (gray) across 3 SCLC cell lines: DMS114, H196, and H446. *P*-values were calculated using a 2-sample *t*-test and were adjusted using the BH method (*P*-adj: DMS114 - 0.00548, H196 - 0.0108, and H446 - 0.925). **(M)** Confocal image of ATF4 immunofluorescence in H446 and H196 +/- ATMi. Nuclei are stained with DAPI. Scalebar = 25 µm. **(N)** H446 and H196 treated with rapamycin (100 nM, 72 h) with or without ATMi (AZD0156), immunoblotted for p-mTOR, p-4EBP1, total-4EBP1, ATF4, and vinculin as loading control. *carcinoid.

For further validation and to examine downstream signaling alterations following ATM inhibition, we conducted RNA sequencing (RNA-seq) in highly susceptible SCLC cell lines (H446, H196, DMS114, H720) pre- and post-treatment with 0.5 µM AZD0156 for 72 hours (**Supplementary Fig. S2B-E**). Two major pathways, mTORC1 signaling and the UPR, were consistently the top suppressed pathways across these cell lines post-ATM inhibition (**Fig. 2B-C**). Specifically, mTORC1 targets were suppressed in H446, H196, and DMS114 (**Fig. 2D**) after pharmacological inhibition of ATM. Western blot analysis in H446 and H196 confirmed that AZD0156 treatment reduced phosphorylated AKT at Ser473 and Thr308, corroborating the CRISPR screen results. Since p-AKT at Thr308 phosphorylates TSC2 at Thr1462 to activate mTORC1, its downregulation suppresses this axis. Consequently, phosphorylated 4EBP1, a downstream effector of mTORC1, was also reduced (**Fig. 2E**) post-AZD0156 treatment.

Prior work established that ATF4, a central TF of UPR and integrated stress response (ISR), is downstream of mTORC1.^29^ The UPR pathway is among the 2 top suppressed pathways from our RNA-seq data (**Fig. 2B**). Consistent with this, ATF4 expression was suppressed following ATM inhibition in both H446 and H196 (**Fig. 2E-F, Supplementary Fig. S2F**. ATF4 and p-4EBP1 downregulation were confirmed using small interfering RNA (siRNA) mediated knockdown of *ATM* in H446 (**Fig. 2F**).

We observe that p-4EBP1 is reduced following ATM inhibition. Unphosphorylated 4EBP1 binds eIF4E and suppresses formation of the eIF4F complex, thereby inhibiting 5ꞌ-cap-dependent translation.^30^ To examine this further, we performed immunoprecipitation using total 4EBP1 and immunoblotted for total eIF4E. ATM inhibition resulted in a significantly increased ratio of 4EBP1 bound to eIF4E in H196 (**Fig. 2G-H**). We next investigated whether reduced 5ꞌ-cap-dependent translation could be functionally validated. Phosphorylated eIF4E, which promotes cap-dependent translation, was downregulated after ATM inhibition (**Fig. 2I**). Since ATF4 is also translated via a cap-dependent mechanism, we hypothesized that suppression of this axis could explain the observed ATF4 downregulation.

To evaluate the relevance of *ATF4* in the clinical SCLC context, we identified the top 50 genes most positively correlated with *ATM* expression in SCLC patients and performed motif enrichment analysis using Enrichr. *ATF4* emerged as the most enriched TF based on both JASPAR and TRANSFAC databases (**Fig. 2J**, **Supplementary Table S2**). SCLC tumors with high tumor mutational burden (TMB) were associated with elevated UPR signaling and worse prognosis.^31^ In line with this, *ATM* expression in SCLC patient samples (*n* = 944) demonstrated strong correlations with both UPR pathway scores (ρ = 0.78) and *ATF4* target gene score (ρ = 0.63), indicating transcriptional alignment between *ATM* expression and UPR signaling (**Fig. 2K, Supplementary Fig. S2G**). Genes scored in **Supplementary Fig. S2G** are ATF4 targets with known involvement in amino acid metabolism, antioxidant pathways, ferroptosis regulation, and UPR, including *ATF4*, *PSAT1*, *GCLC*, *SLC7A11*, *GPX4*, and *SESN2*. We see a coordinated upregulation of *ATF4* target programs and metabolic regulators in ATM-high tumors. RNA-seq confirmed that *ATF4* target genes were significantly downregulated in response to ATM inhibition by AZD0156 (**Fig. 2L**). ATM inhibition also reduced the nuclear localization of ATF4, contributing to its impaired transcriptional activity (**Fig. 2M**).

To probe the extent of UPR inhibition, we examined the levels of canonical UPR effectors including BiP and p-eIF2α, which were also reduced in H446 and H196 alongside ATF4 (**Supplementary Fig. S2F**) upon ATM inhibition. Interestingly, while ATM inhibition suppressed p-PERK, it increased p-GCN2, another eIF2α kinase activated under amino acid deprivation.^32^ Despite this increase in p-GCN2, total p-eIF2α was decreased, prompting investigation of the phosphatase p-PP1α, which was elevated following ATM inhibition (0.5 μM). This suggests a negative feedback loop that suppresses UPR by dephosphorylating eIF2α (**Supplementary Fig. S2F**).

We next wanted to delineate the signaling cascade connecting the AKT pathway to ATF4. Treatment of cells with rapamycin, an mTORC1 inhibitor, resulted in reduced ATF4 expression, which was further decreased when combined with ATM inhibition (**Fig. 2N**). In contrast, genetic knockdown of ATF4 did not alter p-4EBP1 levels (**Supplementary Fig. S2H**), supporting the finding that ATF4 is downstream of mTORC1 signaling. To further evaluate upstream control of this pathway, we treated H446 cells with MK-2206, an AKT inhibitor. Combined treatment with MK-2206 and AZD0156 synergistically reduced both p-4EBP1 and ATF4 levels (**Supplementary Fig. S2I**), reinforcing the presence of an AKT to mTORC1 to ATF4 signaling cascade in ATM-susceptible cells. Collectively, ATM inhibition in SCLC disrupts PI3K-AKT signaling, resulting in mTORC1 and UPR suppression. Disruption of the 5ꞌ-cap-dependent translation through the mTORC1-4EBP1-eIF4E axis causes limited ATF4 translation and nuclear activity. This regulatory cascade underscores ATF4 as a key downstream effector of ATM signaling and suggests that disruption of this axis may compromise stress resilience in SCLC cells. These effects are supported by transcriptomic, proteomic, and patient cohort analyses, identifying ATM as a critical regulator of proteostasis in SCLC.

### ATM inhibition suppresses MYC and ATF4 to disrupt amino acid homeostasis in SCLC

SCLC is characterized by high metabolic demands and dependency on adaptive stress response pathways.^33^ *ATF4* and *MYC* are key regulators of amino acid homeostasis and stress tolerance in cancer cells, operating through transcriptional and translational control of genes involved in proteostasis, redox balance, and biosynthesis.^34,35^ Previous studies have demonstrated that *MYC* and *ATF4* co-regulate several metabolic programs in diverse malignancies, including triple-negative breast cancer, and that their cooperation is associated with poor prognosis.^36–39^ In our analysis of 271 SCLC patients (IMpower133 dataset), high *MYC* target gene expression was associated with worse prognosis (**Fig. 1H**) and was enriched in cell lines highly susceptible to ATM inhibition (**Fig. 1G**).

To establish whether SCLC cells rely on ATF4 and MYC under baseline conditions, we assessed their activity in SCLC models. Assay for transposase-accessible chromatin using sequencing (ATAC-seq) analysis in the ATMi-sensitive H82 cells revealed open chromatin peaks at the *ATF4* gene locus (**Fig. 3A-top panel**), suggesting high transcriptional priming. MYC-ChIP-seq across SCLC lines with varying ATMi sensitivity (SW1271, H128, H2171) showed MYC enrichment at the *ATF4* promoter (**Fig. 3A-bottom panels**), potentially indicating a mode of regulation of *ATF4* transcription by MYC.

**Figure 3.**
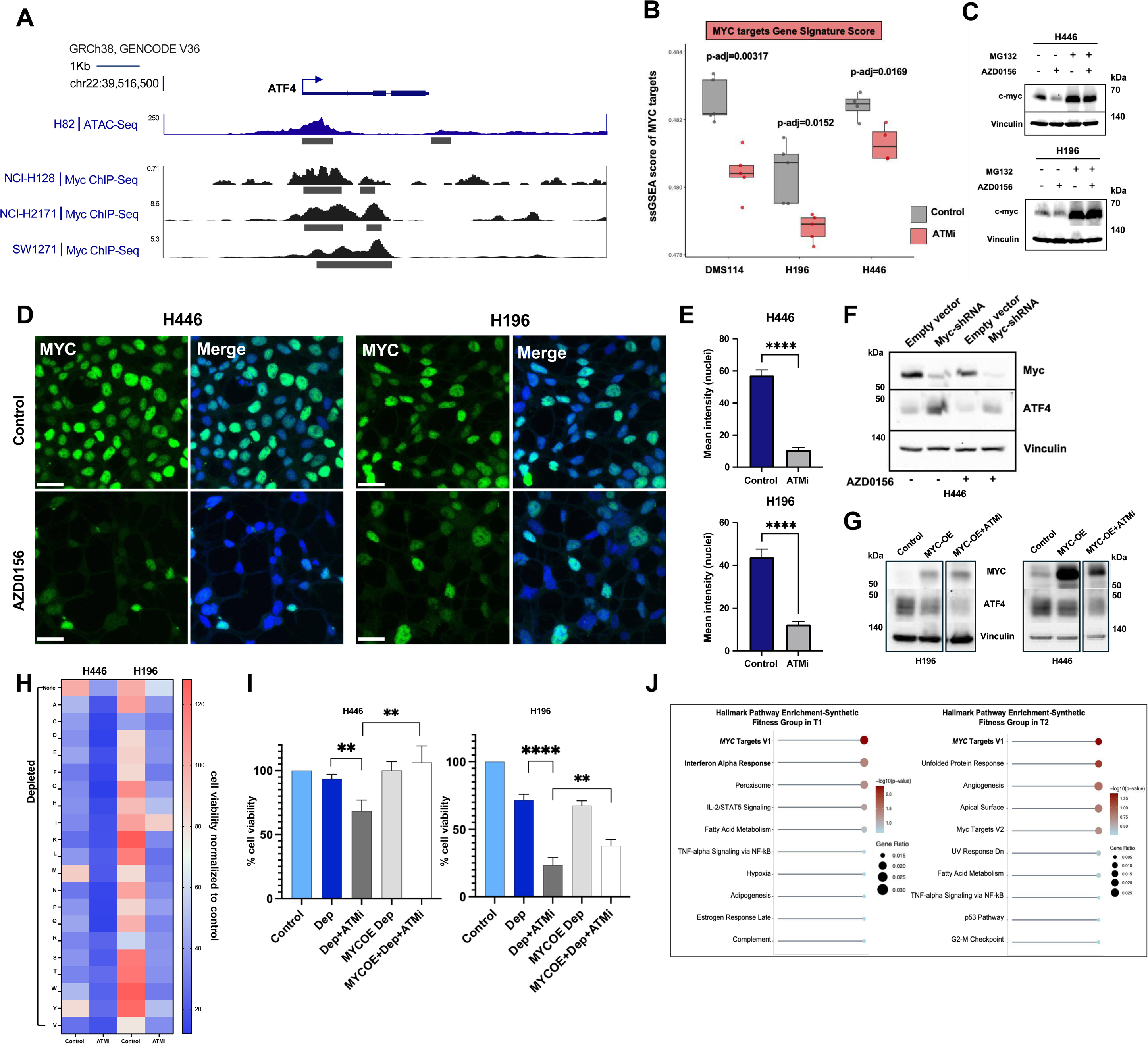
ATM inhibition disrupts a MYC-ATF4 coregulation axis and results in proteotoxic stress. **(A)** ATAC-seq track showing chromatin accessibility at the ATF4 gene locus in H82 SCLC cells. MYC ChIP-seq tracks illustrating MYC binding at the ATF4 promoter region across 3 SCLC cell lines: NCI-H128, NCI-H2171, and SW1271. Gene orientation and structure are displayed using the UCSC Genes track (University of California, Santa Cruz, genome browser), with ATF4 transcribed from left to right on chromosome 22. Blue bars indicate annotated exons. Black bars beneath the ATAC-seq track indicate significant peak calls, marking regions of accessible chromatin, while black bars beneath the ChIP-seq tracks denote significant MYC binding peaks. **(B)** Boxplots showing MYC target gene signature scores in SCLC cells treated with ATMi AZD0156 (red) compared to DMSO controls (gray) across 3 SCLC cell lines: DMS114, H196, and H446. *P*-values were calculated using a 2-sample *t*-test and were adjusted using the BH method (*P*-adj: DMS114 - 0.00317, H196 - 0.0152, and H446 - 0.0169). **(C)** Western blot of H446 and H196 showing expression of MYC after ATM inhibition +/- MG132 (inhibitor of proteosome-mediated degradation). Vinculin is visualized as a loading control. **(D-E)** Confocal image of SCLC models (H446, H196) +/- ATMi showing immunofluorescence for MYC. DAPI was stained to visualize nuclei. Scalebar = 25 µm. Mean intensity of MYC in the nucleus was quantified with or without ATMi. *P*-value was calculated by Student’s *t*-test. **(F-G)** Western blot of H446 cells stably expressing MYC-shRNA **(F)** or in H446 and H196 stably overexpressing MYC **(G)** compared to vector control, +/- ATMi (AZD0156) probed for MYC, ATF4, and vinculin. **(H)** Heatmap showing cell viability (CTG) of H446 and H196 treated with or without ATMi (AZD0156) and depleted of 1 amino acid at a time. **(I)** Bar plot showing cell viability (measured by CTG) in complete and depleted/minimal media +/- ATMi and/or stably overexpressing MYC. *P* values were calculated using Student’s *t*-test. **(J)** Functional enrichment analysis of top-ranked genes identified from the CRISPR screen. Genes selected from the lower end of the sigmoid-ranked essentiality curve were analyzed for pathway enrichment. Lollipop plots display the top enriched biological processes at T1 and T2, with statistical significance indicated by -log₁₀(nominal *P*-value) (color scale) and gene ratio (dot size). Significant pathways are indicated in bold.

In H82, the *MYC* gene locus also shows chromatin accessibility indicating MYC activity in SCLC cells under normal conditions (**Supplementary Fig. S3A**). However, ATM inhibition shows a marked downregulation of MYC activity as depicted by the *MYC* target gene score across 3 ATMi-sensitive SCLC cell lines (H446, H196, and DMS114) (**Fig. 3B**).

We next investigated the effect of ATM inhibition on MYC expression. In H446, H196, and DMS114, AZD0156 treatment reduced both *MYC* mRNA and protein levels (**Fig. 3C, Supplementary Fig. S3B)**. Moreover, treatment with a proteasomal inhibitor (MG132) rescued MYC protein expression, implicating ATM inhibition-mediated proteasomal degradation of MYC in SCLC models (**Fig. 3C**). Finally, nuclear localization of MYC was diminished in H446 and H196 following AZD0156 treatment (**Fig. 3D-E**). MYC is translated via the 5ꞌ-cap-dependent translation machinery, and our previous observation of downregulated p-eIF4E and higher binding of eIF4E by 4EBP1 in the AZD0156-treated condition indicates this to be an active axis of MYC regulation (**Fig. 2G-I**).

To examine the functional interplay between MYC and ATF4, we modulated *MYC* expression genetically. shRNA-mediated or pharmacological knockdown of *MYC* led to an upregulation of ATF4 protein levels (**Fig. 3F, Supplementary Fig. S3C**), while constitutive MYC overexpression in H446 and H196 suppressed ATF4 (**Fig. 3G**). Interestingly, ATF4 overexpression modestly downregulates MYC in both H446 and H196, which is more pronounced on combining ATM inhibition (**Supplementary Fig. S3D**). These findings indicate a bidirectional regulatory balance in which MYC abundance balances ATF4 expression. However, regardless of MYC levels, ATM inhibition further suppressed ATF4 expression in both H446 and H196, consistent with ATM being upstream of this regulatory axis (**Fig. 3G**).

Together, these results define a distinct transcriptional and translational program suppressed by ATM inhibition. ATM inhibition triggers a coordinated downregulation of mTORC1, ATF4, and MYC in SCLC, impairing key regulators of amino acid metabolism and stress adaptation.^29,40,41^ MYC and ATF4 exist in a dynamic regulatory loop, both disrupted by ATM suppression, pointing to a convergent vulnerability in SCLC metabolic control.

### ATM exacerbation impairs amino acid homeostasis in nutrient-limiting conditions and reveals MYC-driven resistance mechanisms

Since ATM inhibition leads to MYC, ATF4, and mTORC1 suppression, the key regulators of metabolic pathways, we next investigated the functional consequences of this transcriptional axis in nutrient-limited environments. Cancer cells frequently rely on de novo amino acid biosynthesis to support proliferation under metabolic stress. We hypothesized that ATM inhibition compromises this compensatory ability in SCLC.

To test this, we conducted a nutrient dropout assay in which individual amino acids were removed from culture media in the presence or absence of ATM inhibition by AZD0156, and cell viability was plotted in a heatmap. Cell viability was measured relative to that in control media without ATMi, which was considered 100% viable. In control conditions, amino acid depletion resulted in moderate reductions in cell viability, reflecting intact compensatory mechanisms via biosynthesis. However, in cells treated with AZD0156, dropout of specific amino acids caused significantly greater reductions in viability in both H446 and H196, indicating impaired metabolic adaptation (**Fig. 3H**). These results suggest that the repression of the ATM-mediated signaling cascade on AZD0156 treatment undermines the cellular capacity to maintain amino acid homeostasis under stress.

From our previous findings, MYC turned out to be a major player of ATM downstream signaling. To determine whether MYC suppression is a key driver of this vulnerability, we examined whether overexpression of MYC could rescue the viability of SCLC cells in nutrient-limited, ATM-inhibited conditions. Indeed, *MYC* overexpression restored cell viability in AZD0156-treated H446 and H196 cultured under nutrient-depleted conditions **(Fig. 3I**), providing direct evidence that MYC restoration counteracts the detrimental effects of ATMi on amino acid stress response.

To further investigate MYC function in early adaptation to ATM inhibition, we analyzed our genome-wide CRISPR-Cas9 knockout screen performed in the presence of AZD0156. *MYC* target genes were the top depleted genes after ATM inhibition, suggesting that MYC inactivation sensitizes cells to ATM inhibition across both T1 and T2 time points (**Fig. 3J, Supplementary Fig. S3E-F**). These findings support a model in which suppression of MYC sensitizes cells to ATMi, but at the same time residual MYC activity becomes essential for cell survival. This dual role highlights a potential synthetic vulnerability that could be leveraged therapeutically.

ATM inhibition disrupts amino acid compensation in nutrient-limiting conditions by impairing MYC and ATF4 function. MYC overexpression rescues this metabolic stress, and MYC targets emerge as early dependencies in genome-wide CRISPR screens under ATMi, highlighting a MYC-driven resistance mechanism and another therapeutic opportunity.

### ATM inhibition disrupts GSH recycling and induces metabolic vulnerabilities in SCLC

We demonstrated that ATM inhibition compromises amino acid homeostasis and proteostasis in SCLC. To further investigate ATM-mediated metabolic rewiring, we performed a targeted metabolomics analysis using a selected central carbon metabolism panel (∼200 metabolites) on 2 representative SCLC cell lines, H446 and H196, pre- and post-AZD0156 treatment. Analysis revealed several metabolites that were elevated upon ATM inhibition in each cell line (**Supplementary Fig. S4A-B**), with 8 metabolites commonly upregulated across both models. These included 5-oxoproline (pyroglutamic acid), creatinine, L-glutamine, guanine, L-methionine, folic acid, and argininosuccinic acid (**Fig. 4A-B**). Among these, 5-oxoproline exhibited high abundance in both H196 and H446 cell lines. This metabolite is a key intermediate in the GSH recycling pathway, also known as the gamma-glutamyl cycle (**Fig. 4C**).^42,43^ Under physiological conditions, 5-oxoproline is converted into glutamate and cysteine, which then contribute to the synthesis of the antioxidant GSH.^42,44^ The accumulation of 5-oxoproline upon ATM inhibition suggested a blockade in the GSH recycling pathway and a potential reduction in cellular GSH. To test this hypothesis, we quantified total GSH and its oxidized form, GSSG, in H196 cells following ATM inhibition with AZD0156, with or without rescue by N-acetyl cysteine (NAC) under both complete and nutrient-depleted media conditions. ATM inhibition caused a significant reduction in GSH levels, accompanied by an increase in GSSG (**Fig. 4D-E**), indicating impaired redox homeostasis. Supplementation with N-acetyl cysteine (NAC), a cell permeable cysteine precursor, counteracted this deficit and effectively restored GSH levels under complete media conditions, but only partially rescued GSH in nutrient-depleted media (**Fig. 4F**). Consistent with this trend, relative GSSG levels followed a similar pattern (**Supplementary Fig. S4C**). Corresponding viability assays demonstrated that NAC fully rescued cell survival in complete media but failed to do so under nutrient-depleted conditions in the context of ATM inhibition (**Fig. 4G**).

**Figure 4.**
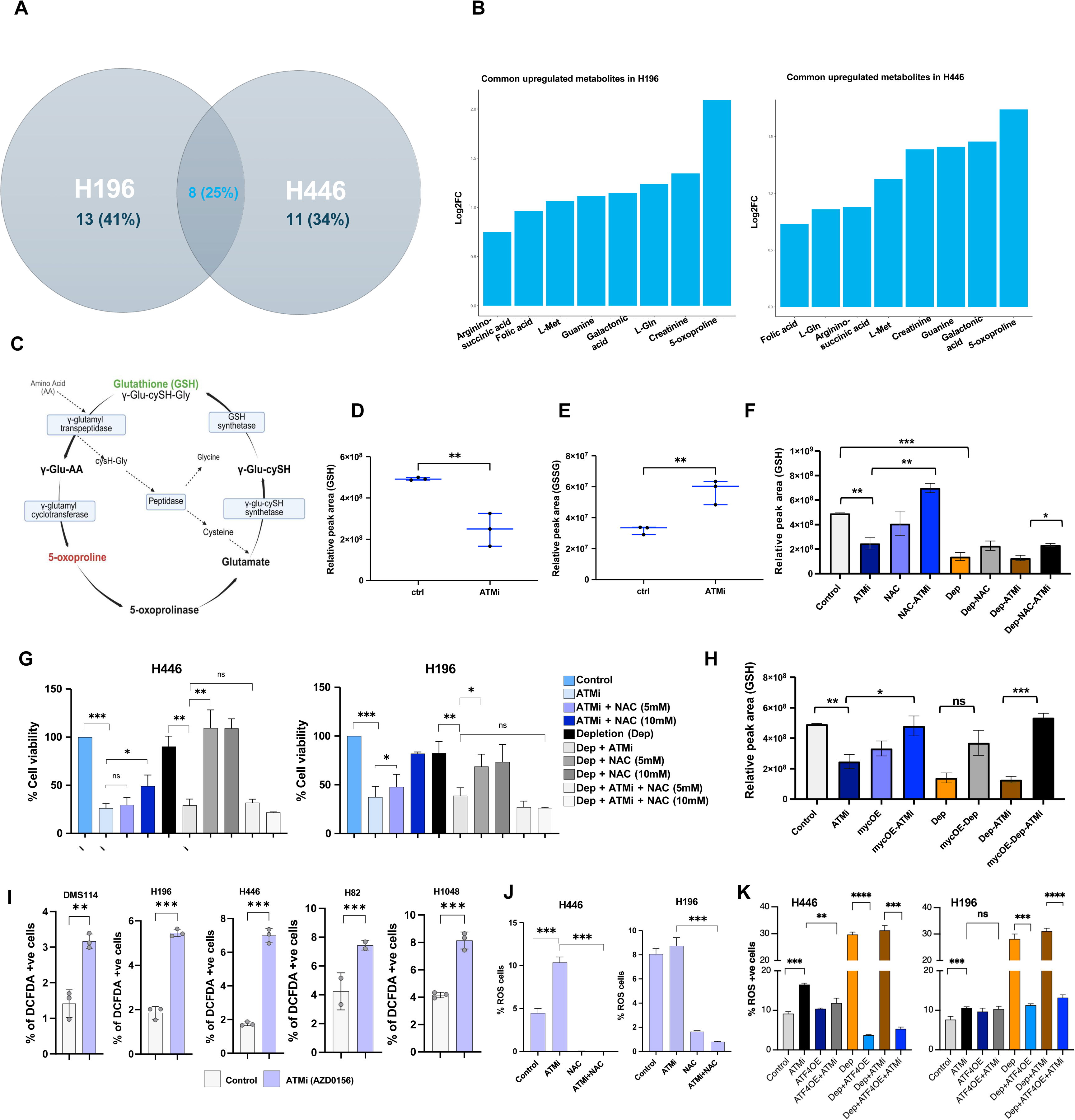
GSH recycling is maintained by functional ATM and preserves redox homeostasis in SCLC. **(A)** Venn diagram showing the number of unique and common upregulated metabolites in H446 and H196 after ATM inhibition **(B)** Bar plot depicting 8 metabolites upregulated in both H446 and H196 cell lines following ATMi. Log 2FC shows the difference between the conditions. **(C)** Schematic representation of the γ-glutamyl GSH cycle. **(D-E)** Dot plots representing the relative abundance of GSH **(D)** and GSSG **(E)** in H196 following ATMi. **(F)** Relative intracellular GSH levels following ATMi +/- NAC in complete or minimal/depleted (dep) media. **(G)** Bar plot showing cell viability of H446 and H196 in complete and depleted/minimal (dep) media +/- ATMi +/- NAC. **(H)** Relative intracellular GSH levels following ATMi and/or stably overexpressing MYC cultured in complete or depleted media. **(I-K)** Relative intracellular ROS in 5 SCLC models **(I)**, +/- ATMi (AZD0156) +/- NAC in H446 and H196 **(J)**, and +/- ATMi and/or transient ATF4 overexpression cultured in complete or depleted media in H446 and H196 **(K)**. Data represent mean ± SEM of *n* = 3 biological replicates. Statistical significance was determined by 1-way Student’s *t*-test (\**P* < 0.05; ** *P* < 0.01; *** *P* < 0.001, **** *P* < 0.0001).

These results indicate that under nutrient limitation, SCLC cells require not only exogenous cysteine but also intact de novo metabolic pathways to sustain GSH biosynthesis. Given that MYC plays a critical role in metabolic adaptation, we tested whether *MYC* overexpression could restore GSH levels in the presence of ATM inhibition. Indeed, *MYC* overexpression reinstated intracellular GSH in both complete and depleted media, confirming the importance of MYC-regulated programs in overcoming ATM-induced metabolic stress (**Fig. 4H, Supplementary Fig. S4D**).

Taken together, these findings suggest that ATM inhibition leads to the accumulation of 5-oxoproline and depletion of intracellular GSH, consistent with impaired glutathione recycling. This disruption compromises antioxidant capacity and cell viability, particularly under nutrient-limited conditions. Rescue experiments support a role for MYC-driven metabolic programs in restoring GSH homeostasis and alleviating ATMi-induced metabolic stress in SCLC models.

### ATM inhibition induces oxidative stress through loss of redox homeostasis and ATF4 downregulation

Since we observed depletion in intracellular GSH post-ATM inhibition, we next examined whether oxidative stress was induced in SCLC cells. We assessed ROS levels using DCFDA staining in 5 SCLC cell lines (H446, H196, H1048, H82, DMS114) post-AZD0156 treatment. All tested models exhibited increased intracellular ROS upon ATM inhibition (**Fig. 4I, Supplementary Fig. S4E-G**). NAC supplementation effectively reduced ATMi-induced ROS levels in complete media (**Fig. 4J, Supplementary Fig. S4H**). However, NAC supplementation failed to provide full rescue to ATMi-induced ROS levels in nutrient-depleted conditions (**Supplementary Fig. S4I**). These findings suggest that under metabolic stress, external cysteine is insufficient to mitigate ROS accumulation without complementary metabolic support.

To better understand this limitation, we examined the role of ATF4, a central transcriptional regulator of redox homeostasis, amino acid metabolism, and proteostasis. ATF4 governs the expression of metabolic enzymes and transporters, such as ASNS, xCT, PSAT1, and CHAC1.^41^ Given its established role in oxidative stress responses, we evaluated whether ATF4 overexpression could alleviate ROS induced by ATM inhibition in nutrient-depleted media (**Fig. 4K**). Expectedly, ATF4 overexpression successfully reduced ROS levels in both H446 and H196 under nutrient-limited conditions post-ATMi treatment, whereas NAC failed to do so. This result indicates that ATF4 downregulation under ATM inhibition disrupts endogenous redox programs, impairing the ability of SCLC cells to synthesize antioxidants such as GSH in the absence of nutrient support. Furthermore, genetic knockdown of *ATF4* using siRNA additively increased ROS under ATMi treatment (**Supplementary Fig. S4J**), further confirming its essential role in maintaining redox balance.

ATM inhibition induces intracellular ROS by disrupting redox homeostasis, an effect that is partially reversible by cysteine supplementation in nutrient-rich environments. However, in nutrient-limited settings, downregulation of *ATF4* prevents metabolic compensation and exacerbates oxidative stress. These findings highlight the critical role of ATF4 in sustaining antioxidant defense under metabolic stress and establish the disrupted ATM-downstream dysregulation as a driver of redox imbalance in SCLC.

### ATM inhibition induces ferroptosis through lipid peroxidation and redox imbalance in SCLC

Given that ATM inhibition induces elevated levels of intracellular ROS, we next investigated whether this redox dysregulation contributes to ferroptosis, a non-apoptotic and iron-dependent form of regulated cell death characterized by GSH depletion and lipid peroxidation. A previous study showed ferroptosis as a vulnerability in SCLC.^45^ Hence, we investigated the level of ferroptosis-related genes in a comprehensive cohort of SCLC clinical samples. In a cohort of 944 SCLC patient samples, expression of a curated anti-ferroptosis-related gene set (including *TFRC, SLC7A11, IREB2, VDAC3, ACSL4, COQ2, GCLC, GPX4, NCOA4, FTH1,* and *AIFM2*) demonstrated a strong positive correlation with *ATM* expression (**Supplementary Fig. S5A**), supporting a transcriptional association between ATM signaling and ferroptosis evasion. Ferroptosis evasion genes, such as *TFRC* and *SLC7A11*, are consistently highly expressed across multiple cancer types and have been proposed as therapeutic targets.^46,47^ Comparative transcriptomic analysis of published datasets confirmed that both *TFRC* and *SLC7A11* were significantly upregulated in SCLC tumors relative to normal tissue (**Fig. 5A-C**).

**Figure 5.**
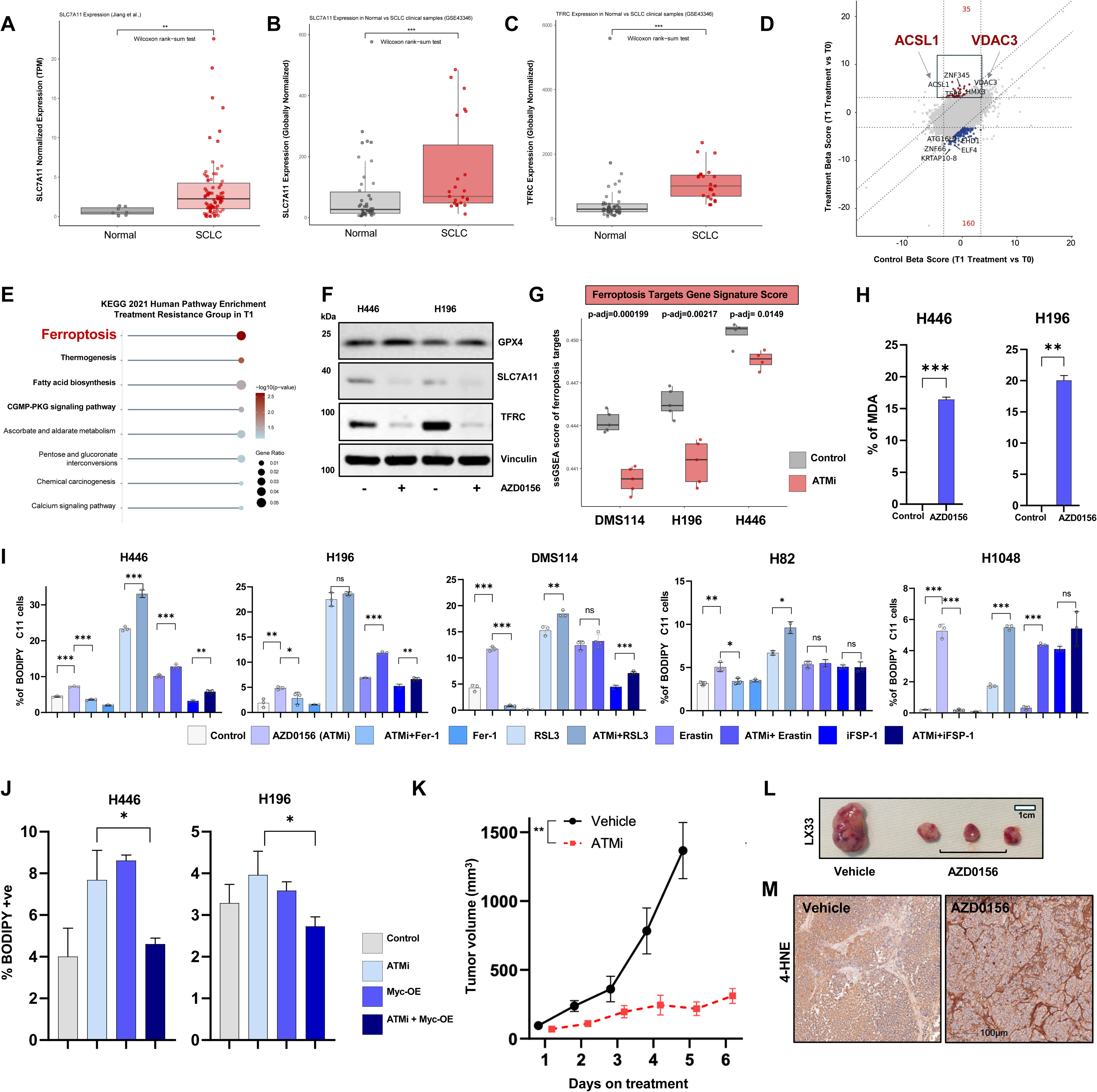
ATM inhibition results in ferroptosis and cell death in SCLC. **(A)** Boxplot representing SLC7A11 mRNA expression from the Jiang et al. microarray dataset of SCLC compared to normal lung tissue (TPM-normalized RNA-seq, Wilcoxon rank-sum test, ** *P* < 0.01). **(B-C)** Boxplot representing SLC7A11 **(B)** and TFRC **(C)** expression in SCLC clinical samples relative to normal controls in the GSE43346 microarray dataset using globally normalized counts (\*\*\**P* < 0.001). **(D)** Nine-square plot comparing gene-level beta scores at timepoint T1 versus T0. The *x*-axis represents gene enrichment or depletion in the control condition, while the *y*-axis shows the same for the treatment condition. Genes in the top-center quadrant (deep-red, *n* = 35) exhibit strong enrichment under treatment but not control, whereas genes in the bottom-center quadrant (blue, *n* = 160) show strong depletion under treatment but not control. Genes in gray represent non-significant hits. Top candidate genes are labeled, including positive hits such as ZNF345, ACS1 and VDAC3, and negative hits such as ATG16L2, EIF4, and ZNF66**. (E)** Functional enrichment analysis of top-ranked genes identified from the CRISPR screen. Genes selected from the upper end (n=35 genes) of the sigmoid-ranked essentiality curve were analyzed for pathway enrichment. Lollipop plot displays the top depleted biological processes at T1, with statistical significance indicated by -log₁₀(*P*-value) (color scale) and gene ratio (dot size). Enriched terms reflect biological processes preferentially required under ATM inhibition (AZD0156). **(F)** Western blot of top ferroptosis markers SLC7A11, TFRC, and GPX4 in whole-cell lysates from H446 and H196 +/- ATMi. Vinculin was visualized as a loading control. **(G)** Boxplots showing ferroptosis gene signature scores in SCLC cells treated with ATMi AZD0156 (red) compared to DMSO controls (gray) across 3 SCLC cell lines: DMS114, H196, and H446. *P*-values were calculated using a 2-sample *t*-test and were adjusted using the BH method (*P*-adj: DMS114 - 0.000199, H196 - 0.00217, and H446 - 0.0149). **(H)** Bar plot representing relative MDA content in whole-cell lysates of H446 and H196 treated with ATMi (AZD0156). **(I)** Bar plot representing relative lipid peroxidation at 24 h measured by BODIPY/C11 positive staining in FACS +/- ATMi (AZD0156) in SCLC models. Ferroptosis inducers (RSL3, iFSP1, erastin) or ferroptosis inhibitors (Fer-1) were used alone or in combination with AZD0156. **(J)** Lipid peroxidation measurement by BODIPY/C11 in H446 and H196 treated with ATMi (AZD0156) with or without stable overexpression of MYC in complete media. Data represent mean ± SEM of *n* = 3 biological replicates. Statistical significance was determined by 1-way Student’s *t*-test (\**P* < 0.05; ** *P* < 0.01; *** *P* < 0.001, \*\*\**\* P* < 0.0001). **(K)** Graph depicts tumor volume of mice over time treated with vehicle or ATMi (AZD0156), with tumor generated subcutaneously from LX33. 0 mg/kg AZD0156 was administered 5/7 days as an intraperitoneal injection. Data are represented as mean ± SEM (*n* = 6 per group). **(L)** Representative image of LX33 tumor sizes from **(K)**. **(M)** IHC images of LX33 tumors +/- AZD0156 probed for expression of 4-HNE, a marker of lipid peroxidation.

Ferroptosis is especially relevant in the context of antioxidant stress, and its initiation is tightly linked to the availability of cysteine and the functionality of the GSH synthesis machinery. The role of ferroptosis in ATMi-mediated response was further highlighted by our genome-wide CRISPR-Cas9 screen of SCLC model H196. The top enriched hits in the early timepoint were involved in ferroptosis, namely *VDAC3* and *ACSL1* (**Fig. 5D-E, Supplementary Fig. S2A**), demonstrating that ferroptosis is necessary for ATMis to kill cells (**Supplementary Fig. S5B-C**). ATM inhibition in SCLC cell lines H446 and H196 led to an appreciable downregulation of TFRC and SLC7A11 (**Fig. 5F**), along with other key ferroptosis-related genes (**Fig. 5G, Supplementary Fig. S5D**). To quantify ferroptosis susceptibility, we applied the FeSig score developed by Zhang et al.,^48^ which reflects transcriptional vulnerability to ferroptosis. High FeSig scores have previously been linked to poor prognosis and resistance to ferroptosis through stromal interactions. ATM inhibition resulted in a significant reduction in the FeSig score in SCLC models, indicating a functional shift toward ferroptotic sensitivity (**Supplementary Fig. S5E-F**).

To assess whether this transcriptional susceptibility translated into lipid peroxidation, we quantified malondialdehyde (MDA) levels, a widely accepted surrogate marker for lipid peroxidation.^49^ MDA levels were significantly increased upon ATM inhibition in both H446 and H196 cells, consistent with the onset of lipid oxidative damage (**Fig. 5H**). Using BODIPY C11 lipid peroxidation analysis, we profiled a panel of SCLC cell lines (H446, H196, H1048, H82, DMS114) treated with ATMi, either alone or in combination with ferroptosis modulators. ATM inhibition alone induced lipid peroxidation across all cell lines and synergistically amplified the effect of ferroptosis inducers erastin and RSL3 (**Fig. 5I**). Moreover, co-treatment with the ferroptosis inhibitor ferrostatin-1 (Fer-1) effectively abrogated ATMi-induced lipid peroxidation in all tested models. These results were corroborated with a second ATMi, KU55933. In H446 and H196 cells, Fer-1 treatment rescued KU55933-induced lipid peroxidation (**Supplementary Fig. S5G**). Together, these findings indicate that ATM inhibition induces ferroptotic stress in SCLC, which can be modulated by pharmacologic agents targeting ferroptotic machinery.

ATM inhibition downregulates key ferroptosis-evasion genes and promotes lipid peroxidation in SCLC cells. These effects are enhanced by ferroptosis inducers and rescued by ferroptosis inhibitors, confirming ferroptosis as a principal mode of cell death driven by ATM inhibition.

### Cysteine limitation and suppression of ATF4 and MYC drive ferroptotic vulnerability upon ATM inhibition

Since ferroptosis is initiated by redox imbalance, we next tested whether antioxidant supplementation could mitigate the effects of ATM inhibition. Treatment with NAC rescued lipid peroxidation in H446 cells under nutrient-deprived conditions and in H196 cells across both nutrient-rich and nutrient-limited environments (**Supplementary Fig. S6A**). These findings underscore the role of ROS in initiating ferroptosis and demonstrate that cysteine supplementation can partially reverse lipid oxidative damage. However, BODIPY staining revealed that lipid peroxidation was noticeable as early as 24 hours following ATMi, highlighting its role as an early and possibly irreversible event in the ferroptotic cascade under continuous ATM inhibition. Since NAC feeds into the γ-glutamyl cycle to enhance GSH synthesis, we hypothesized that cysteine deprivation might synergize with ATM inhibition to exacerbate ferroptotic stress. Indeed, even 12 hours of cysteine withdrawal significantly intensified lipid peroxidation when combined with ATM inhibition, as seen by BODIPY staining (**Supplementary Fig. S6B**), suggesting that cysteine availability is critical for cellular resistance to ferroptotic triggers.

We then focused on ATF4 since it regulates amino acid transport and metabolism, including the expression of SLC7A11, GCLC, and GCLM, all essential for cysteine uptake and GSH biosynthesis. Normally, nuclear accumulation of ATF4 increases during cysteine deprivation, enabling transcriptional compensation.^50–52^ However, under ATM inhibition, the nuclear translocation of ATF4 was impaired in H196, further compromising redox homeostasis (**Supplementary Fig. S6C**). These data highlight a mechanistic role for ATF4 suppression in driving ferroptotic vulnerability upon ATM inhibition.

Finally, we investigated whether restoring MYC, another key component of the ATMi-mediated response, could mitigate lipid peroxidation. Indeed, overexpression of MYC partially reversed lipid peroxidation in both H446 and H196 cells (**Fig. 5J, Supplementary Fig. S6D**). Since MYC is downregulated upon ATM inhibition and plays a central role in redox and metabolic homeostasis, these findings support a model in which coordinated suppression of MYC and ATF4 creates a metabolic bottleneck that renders SCLC cells highly vulnerable to ferroptotic death. Hence, ATM inhibition impairs cysteine metabolism through suppression of ATF4 and MYC, leading to ferroptotic vulnerability. Antioxidant supplementation partially rescues this phenotype, but the early onset of lipid peroxidation and impaired ATF4 signaling under nutrient stress suggests a fundamental disruption of redox homeostasis.

### ATM inhibition induces lipid peroxidation-mediated cell death in patient-derived xenograft in vivo

Our findings so far indicate a ferroptosis-mediated susceptibility to ATM inhibition in SCLC cells in vitro. We next wanted to determine if these ATM inhibition effects could also be observed in vivo. We selected LX33, an SCLC patient derived xenograft model having high expression of NEUROD1, for this part of the study to further unravel the clinical importance of ATM inhibition in SCLC. Assessment of intracellular ROS levels using DCFDA staining in LX33 cells in vitro showed an increase in ROS on ATMi both in complete and depleted growth media conditions (**Supplementary Fig. S6E**). We implanted LX33 subcutaneously in immunocompromised mice and treated them with either vehicle control or AZD0156. Tumor volume measurements showed a significant decrease of 77.26% in tumor volume over time in the AZD0156-treated (50 mg/kg, 5 days a week) group but not in the vehicle group. At day 25, the average tumor volume for the vehicle group was 1,367 mm³, while that for the ATMi-treated group was 310 mm³ (**Fig. 5K-L**). However, the mice remained healthy without any significant differences in body weight during the treatment regime (**Supplementary Fig. S6F**). To confirm the inactivation of ATM in vivo by AZD0156, we stained tissue isolated from vehicle/ATMi-treated mice with a Ser-1981-pATM antibody and found that pATM was only weakly stained in AZD0156-treated tissue as compared to control (**Supplementary Fig. S6G**). This indicated the effective inactivation of ATM in vivo by our AZD0156 treatment routine. We wanted to understand if ferroptosis, which is the major mode of cell death on ATM inhibition in our in vitro studies, is contributing to tumor regression *in vivo*. We looked for the presence of lipid peroxidation within the tumor tissue by staining with 4-hydroxynonenal (4-HNE). 4-HNE is a reactive aldehyde produced during lipid peroxidation of the lipid membrane by excess intracellular ROS. LX33 tumor tissue treated with AZD0156 shows considerably higher levels of 4-HNE as compared to the vehicle (**Fig. 5M**). Together, these data suggest that inhibiting ATM by AZD0156 in vivo can diminish tumor volume due to cell death by ferroptosis without any perceivable toxic effects.

## Discussion

In this study, we uncover a previously unknown role of ATM in SCLC as a central regulator of metabolism. We demonstrate that ATM is highly expressed and activated in SCLC tumors and functionally rewires cellular metabolism and redox balance to promote tumor survival. While ATM is canonically known for its role in DDR, we demonstrate that its inhibition triggers ferroptotic cell death through coordinated suppression of mTORC1, MYC, and ATF4 pathways, highlighting a critical need for redox homeostasis in metabolically stressed SCLC tumors, irrespective of their neuroendocrine or non-neuroendocrine subtype. Multiomic profiling and CRISPR-Cas9 screening approaches used in our study established that inhibition or loss of ATM in SCLC cells attenuates PI3K-AKT-mTORC1 signaling, diminishes cap-dependent translation, and downregulates the oncogenic TF *MYC* and the stress-responsive factor *ATF4*. ATM inhibition affects amino acid uptake and GSH synthesis, driving a collapse of antioxidant defenses and triggering ferroptotic cell death. The ATM-signaling axis also predicts poor prognosis and therapeutic resistance in SCLC. This positions ATM signaling as both a biomarker of aggressive disease and a druggable target whose inhibition affects stress-adaptive programs required for SCLC survival. Our findings advance the role of ATM as a critical regulator of metabolic homeostasis and highlight a previously unknown therapeutic strategy in SCLC.

Ferroptosis-mediated cell death in cancer has been a very promising research avenue, and ferroptosis inducers, such as erastin analogs, RSL3, sulfasalazine, FIN56, and GPX4 inhibitors, have shown potent antitumor activity in preclinical models. However, translating them into safe cancer therapies in the clinic has been difficult. A major hurdle has been systemic toxicity and off-target damage caused by the non-selective nature of ferroptosis induction.^53–56^ An emerging strategy to overcome these limitations is to exploit tumor-specific vulnerabilities, such as oncogene-driven metabolic stress, such that ferroptosis is triggered selectively in cancer cells, sparing normal cells. Inhibiting ATM is a more targeted approach as it affects cancer cells, disabling their ability to manage ROS-induced damage. Normal cells, which are not under such extreme stress, are far less affected by short-term ATM inhibition.^57,58^

We demonstrate that ATM activity sustains the PI3K-AKT-mTORC1 signaling axis in SCLC. Pharmacologic ATM inhibition caused a rapid decrease in AKT phosphorylation and mTORC1 outputs (for example, reduced phosphorylation of 4EBP1), indicating that ATM is required to maintain this pro-survival cascade in SCLC. This was unexpected because previous reports showed that under acute genotoxic or oxidative stress, ATM activates the LKB1-AMPK-TSC2 axis to suppress mTORC1 and induce autophagy as a protective response.^59^ In our SCLC models, ATM appears to play a more permissive role by supporting pro-survival oncogenic signaling pathways like AKT-mTORC1 and maintaining a balance of stress checkpoints. Notably, DMS114 cells, which retain wild-type *RB1* and *PTEN*, differ from canonical SCLC models in their tumor suppressive landscape. Despite this, ATM inhibition in DMS114 leads to similar suppression of mTORC1 signaling and induction of ferroptosis, suggesting that the ATM-mTORC1-ferroptosis axis operates independently of *RB1* or *PTEN* status. This implies that ATM’s role in sustaining redox and metabolic homeostasis may be applicable to all SCLC subgroups, broadening its therapeutic relevance. In hyperproliferative aggressive tumors such as SCLC, cells are known to generate excessive ROS yet evade oxidative cell death by realigning their redox balance through the upregulation of antioxidant systems, such as γ-glutamyl-based GSH synthesis, thioredoxins, and NADPH-generating pathways. Consistently we observed elevated mitochondrial ROS upon ATM inhibition, as evidenced by MitoSOX staining, further underscoring the disruption of redox homeostasis. Intracellular antioxidant systems aid in sustaining intracellular ROS while avoiding thresholds that would otherwise trigger cell-death pathways. These metabolic balances not only contribute to initiation, progression, metastasis, and recurrence, but might also contribute to the inter-tumor heterogeneity of SCLC.^60^

In our study, γ-glutamyl-GSH recycling was found to be compromised on ATM inhibition, likely due to a disruption in amino acid homeostasis and perturbation of the ATM-MYC-ATF4 circuit. ATF4 is a known transcriptional activator of cystine import and GSH biosynthesis pathways.^35^ MYC can also influence glutamine and cysteine metabolism in cancer cells. GCLC, the catalytic subunit of glutamate-cysteine ligase that synthesizes GSH from cysteine and glutamate, is directly regulated by MYC. Thus, inhibition of both ATF4 and MYC in SCLC impairs the import or synthesis of sufficient precursors to maintain GSH levels and cannot sustain expression of antioxidant enzymes. This is in agreement with our data that shows that *MYC* overexpression can completely restore GSH levels, whereas, in ATM-inhibited conditions, cysteine supplementation by NAC is only partially beneficial. Historically, cysteine deprivation was one of the first conditions to induce ferroptosis.^49^ Cysteine is usually imported via the SLC7A11/xCT system and is indispensable for GSH synthesis. When cystine is scarce, GSH levels drop, and cells become vulnerable to oxidative damage and turn on adaptive programs like ISR. Normally, nutrient stress activates GCN2/ATF4 signaling to upregulate amino acid transporters, transsulfuration enzymes, and other compensatory mechanisms.^38^ In our experiments, however, ATM-deficient SCLC cells were unable to mount an effective ATF4-mediated response. As a result, the combination of ATMi and low/minimal nutrients severely compromised cell viability. Since ATF4 and MYC are both critical modulators of the de novo metabolism ^37,61–63^ of amino acids and can regulate the transcriptional output of several enzymes and solute carriers, cells depend on them as rheostats for nutrient availability. ATM inhibition downregulates both ATF4 and MYC, exacerbating nutrient stress. This synthetic lethal interaction between ATM loss and nutrient stress aligns with the idea that the tumor microenvironment, which is often nutrient poor, could unmask latent vulnerabilities in cancer cells. ATM-high SCLC tumors may survive in part by better adapting to fluctuating nutrient and oxygen levels, an ability conferred by ATM’s support of ATF4/MYC and mTORC1 activity. Removing ATM removes this safety net, particularly under microenvironmental stress induced by conventional therapies, leading to ferroptosis and rapid cell death.

We demonstrate that inhibiting ATM downregulates global cap-dependent translation. mTORC1 usually phosphorylates 4EBP1 to liberate eIF4E and drive translation initiation.^64^ In the absence of this inhibitory phosphorylation, 4EBP1 increasingly binds eIF4E. ATMi-treated cells showed accumulation of hypophosphorylated 4EBP1 (the active, translation-repressive form) and eIF4E, indicating reduced cap-dependent protein synthesis. Importantly, among the most translation-sensitive proteins are short-lived oncogenic regulators like MYC and ATF4. We observed that ATM inhibition led to decreased MYC and ATF4 protein levels, even when transcript levels did not fully collapse, suggesting a translational and post-translational effect. A striking effect of ATM inhibition is the dual suppression of MYC and ATF4. Even though it was previously reported^38,39,65^ that cells leverage MYC and ATF4 through a connected axis, a critical balance between the MYC-ATF4 stoichiometry is part of its adaptive resilience. Hence, SCLC cells intrinsically maintain an equilibrium between MYC and ATF4 in a pro-survival manner; ATM inhibition disrupts this balance, leading to detrimental effects on cell viability. MYC has been identified as a biomarker in SCLC, but there are no existing therapies that efficiently target it.^66,67^ Previous studies have identified MYC expression as a key determinant of a cell’s metabolic phenotype, with high *MYC* levels promoting a Warburg-like glycolytic program, while low *MYC* expression is associated with greater reliance on mitochondrial respiration.^67^ ATM inhibition directly targets the metabolic landscape of SCLC cells via ISR cascade involving both ATF4 and MYC.

We found that low-dose ferroptosis inducers (erastin, RSL3) strongly boosted the effect of ATM inhibition, consistent with lowering the threshold for lipid peroxidation-mediated cell death. This approach is in alignment with a recent study showing that mTORC1 inhibitors synergize with ferroptosis inducers to suppress tumor growth in vivo.^58^ In SCLC, ATM inhibition serves as an mTORC1 inhibitor surrogate and combines additional downstream impacts; hence, combining it with ferroptotic stressors or inhibitors of cystine uptake could yield potent antitumor effects.

In summary, this study provides a foundation for a new synthetic lethal approach to SCLC, targeting the Achilles’ heel of metabolic and redox homeostasis via ATM. Here, we report the exploitation of UPR as a major mode of adaptation in SCLC, which is regulated by DDR kinase ATM, through a complex signaling network. This may further account for the heightened dependence of *p53* and *Rb1* loss-of-function (LOF) cancers on major DDR pathways. Our findings reveal that the 2 pivotal determinants of metabolism, *ATF4* and *MYC*, maintain metabolic rhythm in the cell through a tightly regulated and fragile balance. ATM is a key stabilizing force in this system, and its inhibition precipitates into a breakdown of proteostasis. This, in turn, compromises the redox buffering capacity of the cell, culminating in lipid peroxidation-driven ferroptotic cell death. This work identifies ATM as a druggable regulator in SCLC whose inhibition can be utilized in indirect disruption of critical adaptive nodes, including hard-to-target AKT, mTORC1, and MYC, in a single shot. Our in vivo studies support tolerability for ATM inhibition with no detectable systemic toxicity. The accumulation of 4-HNE in tumors upon ATM inhibition displays marked lipid peroxidation, providing in vivo evidence of ferroptosis. From a clinical perspective, these findings highlight ATM as an attractive therapeutic target in SCLC and open the avenue for rational combination strategies. By simultaneously incapacitating DNA repair and metabolic rescue pathways, such combinations could achieve what the standard of care often cannot: the induction of a non-apoptotic form of cell death that SCLC cells are less equipped to escape.

### Limitations of the study

While the current study mainly focused on cytosolic antioxidant GSH recycling, the ATM-dependent regulation of mitochondrial antioxidant defenses represents an important avenue for future investigation. Ferroptotic cells release damage-associated molecular patterns (DAMPs) and lipid mediators that can stimulate immune responses.^68,69^ We observe a remarkable increase in the lipid oxidation product 4-HNE in LX33 mice tumor tissue consistently treated with AZD0156. ATMis not only kill SCLC cells directly but may also heighten their visibility to the immune system, perhaps improving responses to checkpoint blockade. While the current work focuses on ATM-mediated cell-intrinsic metabolic rewiring and antitumor responses in SCLC, further work in immunocompetent models would be interesting to test the antitumor immune response of ATM inhibition. It is yet to be fully understood how ATM inhibition leads to MYC downregulation and enhanced MYC degradation. One plausible mechanism involves the suppression of PI3K-AKT signaling, which is known to stabilize MYC by preventing GSK3β-mediated phosphorylation.^70^ Finally, MYC transcriptionally regulates ATF4, and perturbation of this axis impairs metabolism. Future studies are warranted to understand if *MYC* acts as a repressor of *ATF4* or if there are additional mechanisms involved to maintain homeostasis in the collective stoichiometry of ATF4 and MYC within the cell.

## Resource availability

### Lead contact

Requests for further information and resources should be directed to and will be fulfilled by the lead contact, Triparna Sen.

### Materials availability

This study did not generate any unique reagents or materials as all the materials that were used are commercially available or previously published.

### Data and code availability

For clinical samples from Caris Life Sciences and deidentified sequencing data owned by Caris Life Sciences, qualified researchers can apply for access by contacting A.E. and signing a data usage agreement. RNA-seq and CRISPR screen data have been deposited in the NCBI Gene Expression Omnibus under the accession number GSE307086.

## STAR Methods

### EXPERIMENTAL MODEL AND STUDY PARTICIPANT DETAILS

#### Clinical samples cohort

Comprehensive molecular profiling was performed on real-world tumor specimens from 179,633 patients, submitted to a Clinical Laboratory Improvement Amendments (CLIA)-certified laboratory (Caris Life Sciences, Phoenix, AZ). The study adhered to the ethical principles outlined in the Declaration of Helsinki, the Belmont Report, and the US Common Rule, and was conducted under the exemption criteria outlined in 45 CFR 46.101(b). As the analysis was retrospective and utilized deidentified clinical data, informed consent was not required. Statistical analyses were conducted using JMP software (version 13.2.1, SAS Institute).

### METHOD DETAILS

#### Gene expression profiling for SCLC subtyping

To classify the patient cohort (*N* = 944) into SCLC subtypes, RNA expression data (TPM) for the established and candidate lineage-defining transcription factors ASCL1 (A), NEUROD1 (N), POU2F3 (P), and YAP1 (Y) were converted to *Z*-scores. Samples showing a positive *Z*-score for only 1 of the 4 factors were assigned to the corresponding subtype. Cases with more than 1 positive *Z*-score were categorized as mixed, whereas those with negative *Z*-scores across all 4 factors were designated as transcription factor-negative (TF-Neg). All statistical values are mentioned in Supplementary table S1.

#### IHC and whole tissue slide imaging

Formalin-fixed paraffin-embedded (FFPE) 3-μm sections from archival tissue at initial diagnosis were used for IHC staining. IHC was performed using the Ventana Discovery Ultra (Roche Diagnostics) on RUO Discovery Multimer V2 (v0. 00.0083) platform. This system allows automated baking, deparaffinization, and cell conditioning. All primary antibodies were incubated for 60 minutes at 37 °C. Anti-ATM mouse monoclonal antibody [2C1 (1A1)] (abcam ab78) (1:100X), anti-ATM (phospho S1981) rabbit monoclonal antibody [EP1890Y] (abcam ab81292) (1:50X), and anti-4 hydroxynonenal antibody [HNEJ-2] mouse monoclonal antibody (abcam ab48506) (1:200X) were used as primary for staining. Discovery OmniMap anti-rabbit HRP (760-4311) and anti-mouse HRP (760-4310) were used as secondary antibodies. Discovery ChromoMap DAB kit, Roche Diagnostics (760-159) was used as a DAB detection kit. Mayer hematoxylin was used for nuclear counterstaining. Whole tissue sections on the slide were converted into high-resolution digital data using a NanoZoomer S210 digital slide scanner (Hamamatsu).

#### Cell culture

SCLC cell lines were obtained from the American Type Culture Collection (ATCC) and the European Collection of Authenticated Cell Cultures (ECACC). Cells were cultured in Roswell Park Memorial Institute (RPMI) 1640 medium (Gibco, A10491-01) supplemented with 10% fetal bovine serum, 100 U/mL penicillin, 50 µg/mL streptomycin, and 2 mM L-glutamine. Cultures were maintained at 37 °C in a humidified atmosphere with 5% CO₂ and over 90% relative humidity. All cell lines underwent routine mycoplasma testing and were authenticated by short tandem repeat (STR) DNA profiling prior to use. For experimental consistency, cells were typically used within 2-8 passages after thawing. Cells were stored in liquid nitrogen, in cryovials (Corning) using Bambanker freezing medium (Fisher Scientific, BB01).

#### Drugs used in this study

AZD0156 (Selleckchem, S8375), KU55933 (Selleckchem, S1092), and AZD5305 (Selleckchem, S9875) were obtained from Selleckchem. Other inhibitors used were AKT inhibitor MK-2206 (Medchemexpress), mTORC1 inhibitor rapamycin (Thermo Scientific), apoptosis inhibitor ZVAD (Selleckchem, 187389-52-2), apoptosis inducer (PKC inhibitor) Staurosporin (Selleckchem, NC1828540), ferroptosis inhibitor ferrostatin (Cayman Chemical), xCT inhibitor erastin (Selleckchem, 50-136-4551), GSH peroxidase 4 (GPX4) inhibitor RSL3, and AIFM2/FSP1 inhibitor iFSP1 (Fisher Scientific), MG132 (Cayman Chemical, 133407-82-6), and MYC inhibitor (MYCi975, Selleck Chem, S8906).

#### Cell viability assay

SCLC cell lines were seeded in 96-well plates at a density of 2,000 cells per well and incubated overnight in complete RPMI-1640 medium supplemented with 10% fetal bovine serum, 100 U/mL penicillin, 50 µg/mL streptomycin, and 2 mM L-glutamine. Cells were then treated with either DMSO (vehicle control) or serial concentrations of AZD0156 or KU55933 for 120 hours (5 days). Viability was assessed using the CellTiter-Glo (CTG) Luminescent Cell Viability Assay (Promega, Madison, WI), following the manufacturer’s instructions. IC₅₀ values were calculated using nonlinear regression analysis in GraphPad Prism version 9.0 (GraphPad Software, San Diego, CA).

#### Depletion assays

For the amino acid depletion assay, 3,000 cells were seeded in complete growth media in 96-well plates (Corning, 353296) for CTG analysis and allowed to adhere overnight. The next day, cells were switched to amino acid–free, glucose-free media (MyBioSource, MBS652918) supplemented with all amino acids (1 mM each) except one, generating 20 individual conditions in which a single amino acid was omitted per medium. Amino acids were purchased from Sigma and stock solutions were prepared freshly on the day of the experiment. Glucose (2 g/L) and 10% dialyzed FBS (Thermo Fisher, A3382001) were additionally provided to the culture. Cells were incubated in a humidified incubator at 37 °C for 72 hours with DMSO (vehicle) or AZD0156 added to the media. Cell viability was measured by CTG and analyzed relative to cells grown in complete media containing all 20 amino acids using Graphpad Prism. Depletion/minimal media was prepared using MEM media (Thermo Fisher), 10% dialyzed FBS, 2 mM Glutamax (Thermo Fisher), 17 mM glucose (Sigma), and 1% MEM-vitamin (Thermo Fisher Scientific). Freshly prepared 5-10 mM N-acetyl cysteine (NAC, Sigma) was used for supplementation in ROS-rescue experiments. For cysteine-depletion experiments, cells were cultured in glutamine- and cysteine-free medium (MP Biomedicals, 091646454) supplemented with penicillin-streptomycin and 10% FBS for 12-72 h. All media were equipped with 1% Pen-Strep (Thermo Fisher).

#### *In vivo* ATM inhibitor preparation

For the in vivo experiment, the formulation was prepared as per the manufacturer’s instructions. The ATMi AZD0156 (Selleckchem, S8375) was used with a formulation of 40% propylene glycol, 5% DMSO, 5% Tween 80, and 50% water. Vehicle formulation was used with a solution of 40% propylene glycol, 5% DMSO, 5% Tween 80, and 50% water. At first, the drug was measured in an appropriate amount and dissolved in 5% equivalent DMSO solution. Then, a 40% equivalent volume of PEG (polyethylene glycol) was added and vortexed vigorously for 2-5 min, after which the formulation was mixed and clarified. 5% equivalent Tween 80 was added, and the step of mixing and clarifying was repeated. In this step, some of the drug particles may remain undissolved, and physical methods such as mild sonication and a hot water bath can be used. After that, 50% sterile water was added to the formulation to achieve the final drug formulation. For preparation of the drug formulation, solvents were added sequentially in the order described above. The clarity of the solution obtained after each addition was verified prior to introducing the next solvent.

#### Immunodeficient mouse model and in vivo treatments

A total of 2 × 10⁶ SCLC cells (H446, H196, or PDX-derived LX33) were suspended in a 1:1 ratio of phosphate-buffered saline (PBS) and Matrigel (Fisher Scientific, CB40234) and injected subcutaneously into the flanks of 6-week-old female nude mice (ENVIGO, Boyertown, PA). Once tumors reached an average volume of 100-150 mm³, mice were randomized into vehicle and treatment groups (*n* ≥ 8). AZD0156 was administered intraperitoneally at 50 mg/kg, 5 times per week (5/7 days), while control mice received vehicle (solvent, subcutaneously). For animal experiments, comparisons between 2 groups are done using an unpaired student’s *t*-test. No post-hoc test was performed as we had only 2 comparison groups.

#### Animal studies and tumor monitoring

All animal procedures were conducted in accordance with protocols approved by the Institutional Animal Care and Use Committee (IACUC). Tumor dimensions were measured twice weekly with calipers, and volumes were calculated using the formula: (width² × length × 0.5). Body weights were also monitored biweekly. Humane endpoints included tumor volume exceeding 2 cm in any dimension, volume surpassing 10% of body weight, sustained weight loss (> 10%), or signs of ulceration. Tumor growth kinetics were analyzed using linear mixed-effects regression models to evaluate treatment response.

#### Flow cytometry analysis

Flow cytometry was conducted following standard procedures. Data acquisition was performed using FACSymphony A5 SE in FACSDiva software. Acquired data was further analyzed in FlowJo (v10.6; BD Biosciences, Franklin Lakes, NJ). For cell-cycle analysis, cells were rinsed once in ice-cold PBS, fixed overnight at -20 °C in 70% ice-cold ethanol, and washed again in PBS the following day. Cells were then stained with FxCycle™ PI/RNase solution (Invitrogen) for 15 min at room temperature and analyzed on the cytometer. For apoptosis analysis, 0.5 × 10⁶ cells were seeded in 6-well plates in RPMI medium supplemented with 10% FBS and allowed to adhere overnight. On day 2, cells were treated with the indicated drugs for the specified duration. Following treatment, cells were harvested by trypsinization, washed twice with ice-cold PBS, and once with cold Annexin V binding buffer (BioLegend, 6410914). Cells were then resuspended in 100 µL binding buffer containing 5 µL Annexin V-FITC (BioLegend, 640906) and incubated for 10 min at room temperature in the dark. Propidium iodide (PI) (10 µL) was added, and samples were gently mixed and incubated for an additional 15 min in the dark. Prior to acquisition, 400 µL of binding buffer was added. For data acquisition, negative controls were used for gating. Cells treated with 1 µM staurosporine for 8 h served as positive controls for Annexin V-FITC and PI staining.

For performing FACS-based detection of intracellular ROS or lipid peroxidation, DCFDA and BODIPY/C11 staining protocol were employed, respectively. Cells were seeded either in a 6-well plate (0.2 ×10^6^ cells/well) or in a 12-well plate (0.1 × 10^6^ cells/well) and incubated overnight. Post-seeding, cells were treated with indicated drug(s) or vehicle (DMSO) and incubated at 37 °C for 24 h for BODIPY/C11 assay and 24-72 h for DCFDA ROS assay, respectively. Following the completion of incubation, media was supplemented with 3 μM of C11-BODIPY 581/591 Lipid Peroxidation Sensor (Invitrogen, D3861), and the assay was performed using the previously published protocol.^1^ As RSL3 is a ferroptosis inducer, cells treated with RSL3 for 3 h were used as a positive control for gating high C11/BODIPY stained-cells induced by RSL3 treatment. For DCFDA assay, media was removed completely, followed by PBS wash and treatment with 10 µM DCFDA in PBS, for 30 min at 37 °C before subjecting the suspension to FACS analysis. Vehicle-treated cells were compared to drug-treated cells with a shift in the curve to the right, indicating increased intracellular ROS or lipid peroxidation for DCFDA and BODIPY, respectively. Data was plotted using Graphpad Prism software. An unpaired Student’s *t*-test was used to determine pairwise significance.

#### MitoSOX

The measurement of mitochondrial ROS production was conducted using the MitoSOX™ Red mitochondrial superoxide indicator (Invitrogen). SCLC cells (0.2 ×10^6^) were seeded in 12-well plates; after the respective treatment, the media was removed, and the 5-μM MitoSOX Red in PBS was added and incubated for 20-25 min at 37 °C in the dark. Cells were then trypsinized and washed with PBS, and samples were collected in PBS. The data acquisition process was carried out with thoroughness and precision using a FACSymphony A5 instrument. The comparison between vehicle-treated cells and drug-treated cells was made with a shift in the curve to the right side, indicating increased mitochondrial ROS. The acquired data were further analyzed using FlowJo (v10.6; BD Biosciences, Franklin Lakes, NJ) software. Data was plotted using Graphpad Prism software. An unpaired Student’s *t*-test was used to determine pairwise significance.

#### CRISPR library preparation and data analysis

Genome-wide knockout screening was performed using the Toronto KnockOut v3 (TKOv3) pooled sgRNA library strictly following the Moffat laboratory protocol (Hart et al., 2015; Moffat Lab, University of Toronto). H196 cells were transduced in triplicate at an MOI of 0.35 in the presence of 8 µg/mL polybrene, followed by puromycin selection (3 µg/mL) for 48 h. Cells were maintained at ≥ 200-fold library coverage and cultured for 2-3 population doublings under 2 parallel conditions: control (DMSO) and AZD0156 treatment. At the experimental endpoint, cell pellets were collected and stored at -80 °C for genomic DNA extraction and next-generation sequencing-based sgRNA abundance quantification. Genomic DNA was extracted from cell pellets using the Wizard Genomic DNA Purification Kit (Promega) according to the Moffat laboratory TKOv3 protocol. sgRNA cassettes were amplified by a 2-step PCR to incorporate Illumina TruSeq adapters with unique i5/i7 indices. Purified libraries were quantified by Qubit and sequenced on an Illumina HiSeq platform using single-end 50 bp reads. Schematic diagrams illustrating the experimental workflow were created using BioRender (www.biorender.com) under an institutional license.

#### Western blot

Protein extraction was carried out following previously reported methods^2^ with minor modifications. In brief, cells were harvested in ice-cold PBS and centrifuged at 2,500 rpm for 3 min. The resulting pellet was resuspended in RIPA buffer (Thermo Fisher, 89901) supplemented with sodium fluoride and a phosphatase inhibitor cocktail (Thermo Fisher, 78446). Lysates were kept on ice for 1 h, with gentle vortexing every 5 min, and subsequently centrifuged at 12,000 rpm for 20 min at 4 °C. The supernatant, representing total protein, was collected and quantified using the Coomassie Protein Assay (Thermo Scientific, 1856209). Equal amounts of protein (50 µg) were mixed with Laemmli Sample Buffer (Bio-Rad, 1610747), separated on 10-15% SDS-PAGE gels, and transferred by wet blotting onto 0.45-µm Immobilon-FL PVDF membranes (Millipore, IPFL00010). Membranes were blocked for 1 h at room temperature in Blocker Casein (Thermo Scientific, 37532) prepared in TBS containing 0.1% Tween 20 (Fisher Bioreagents, BP337-500), then incubated overnight at 4 °C with primary antibodies (1:1000 dilution in 5% bovine serum albumin [BSA]). Following TBST washes, membranes were probed with horseradish peroxidase-conjugated secondary antibodies (1:8000 dilution) for 1 h at room temperature. Detection was performed using the SuperSignal™ West Pico PLUS Chemiluminescent Substrate (Thermo Scientific, 34577), and images were captured on an iBright Western Blot Imaging System (Thermo Fisher). All primary antibodies are listed in **Key Resource Table**.

#### Immunofluorescence

Cells were cultured on chamber slides (BD Falcon) with appropriate treatment and then fixed with 4% paraformaldehyde (Thermo Fisher) for 30 min at room temperature. Following permeabilization with 0.1% Triton X-100 (Thermo Fisher) for 8-10 min, slides were rinsed once with PBS and blocked with blocking solution (Thermo Fisher, NC1584859). Primary antibodies, diluted as per manufacturer’s recommendations, were applied overnight at 4 °C. The next day, slides were washed 3 times with PBS and incubated in the dark with Alexa Fluor 488 and/or 594- conjugated secondary antibodies (Thermo Fisher) for 30-60 min. After 3 additional PBS washes, chambers were removed, slides were air-dried briefly, and coverslips were mounted with DAPI-containing mounting medium (Invitrogen, 50-112-8966). Images were captured on a Leica TCS SP8 confocal microscope using Leica LAS X software. All analyses were done using ImageJ-FIJI software.

#### Immunoprecipitation

Cells were lysed in Pierce™ IP Lysis Buffer (Thermo Scientific) following the manufacturer’s protocol for adherent cells. Clarified lysates were incubated with primary antibody and Protein A/G Magnetic Beads (Thermo Scientific™ Pierce™, Fisher PI88803) with end-over-end rotation at 4 °C overnight. Beads were captured on an Invitrogen™ DynaMag™-2 magnet and washed in TBS buffer according to the manufacturer’s instructions. Bound complexes were eluted with Pierce™ IgG Elution Buffer (pH 2.0) and immediately neutralized with Tris-HCl (pH 8.0). Where indicated, input and eluted fractions were analyzed by SDS-PAGE and immunoblotting.

#### Transfection of siRNAs

For siRNA transfection, 0.5 × 10⁶ cells were seeded in 6-well plates and grown to approximately 75% confluence. Cells were transfected with 10 nmol/L anti-ATF4 siRNA (Thermo, AM16708, ID: 122168,122287) or ATM Mission esiRNA pool (Sigma Millipore, EHU089521) or scrambled control siRNA (AM4611, Invitrogen) using lipofectamine RNAiMAX reagent (Thermo Fisher Scientific, 13778030). After 16-20 h of incubation at 37 °C in 5% CO₂, the medium was topped with fresh RPMI to minimize cytotoxicity. Cells were harvested 48 h post-transfection for Western blot analysis to confirm transfection efficiency and for downstream experiments.

#### Lentiviral generation and transduction

Lentiviral particles were generated by co-transfecting 293T cells with the lentiviral backbone construct together with the packaging plasmids psPAX2 and pMD2.G, using lipofectamine LTX reagent and PLUS reagent (Invitrogen, 15338030). Sixteen to 18 h post-transfection, the culture medium was replaced with BSA-enriched DMEM. Viral supernatants were harvested 72 h after transfection, centrifuged to remove cellular debris, and aliquoted for storage at -80 °C. On day 0, SCLC cells were seeded at 1 × 10⁶ per 10-cm dish and allowed to adhere overnight. The following day, culture medium was replaced with 10 mL of fresh RPMI containing 10% FBS and 8 μg/mL polybrene (Tocris, R&D Systems, 7711). Lentiviral stocks stored at -80 °C were rapidly thawed in a 37 °C water bath and placed on ice prior to use. Virus was added to the cells dropwise at a volume determined by prior titration, and plates were incubated overnight at 37 °C. On day 2, the lentiviral-containing medium was replaced with fresh RPMI supplemented with 10% FBS. Puromycin selection (3 µg/mL) was initiated on day 3 and continued for 48 h. MYC shRNA construct ref. seq. NM_002467 (Target seq: CCTGAGACAGATCAGCAACAA) was obtained from Sigma-Aldrich and transfected into SCLC cell lines using lentiviral vector according to the manufacturer’s instructions. Other lentiviral particles used for the study were generated from ATF4-OE construct (Addgene, pRK-ATF4, 26114), or MYC-OE construct (Addgene, MYC- PGK-blast, 190618) using the same protocol.

#### Metabolomics profiling

Polar metabolite profiling of SCLC cells treated with 250 nM AZD0156 or DMSO (vehicle) for 72 h was performed using a 200-metabolite panel at the Metabolomics Core, Icahn School of Medicine at Mount Sinai. Metabolism was quenched by aspirating medium and adding 1 mL of ice-cold 80:20 methanol:water. After overnight incubation at -80 °C, cells were collected and centrifuged (20,000 g, 20 min, 4 °C). Supernatants were dried under vacuum (Genevac EZ-2 Plus, 3 h), resuspended in 30 μL water, vortexed, incubated on ice (20 min), and clarified by centrifugation. Analysis was performed with an Agilent 1290 Infinity II HPLC coupled to a 6495D triple quadrupole MS (negative ESI) using a ZORBAX RRHD Extend-C18 column (2.1 × 150 mm, 1.8 µm). Mobile phases were 3% methanol in water (A) and 100% methanol (B), both containing 10 mM tributylamine, 15 mM acetic acid, and 5 µM medronic acid. The LC gradient (0-40 min) ramped from 100% A to 99% B, followed by acetonitrile backflush and re-equilibration. Flow rate was 0.25 mL/min and column temperature was 35 °C. MRM transitions are listed in **Supplementary Table S3**. Compound identity was confirmed with pure chemical standards. Data analysis (peak integration and signal extraction) was performed with Skyline (v24.1.1.398).

#### Targeted GSH and GSSG panel

Extraction was performed as above, except dried samples were reconstituted in 100 μL of 50% acetonitrile. Metabolites were analyzed by HILIC on a Vanquish Horizon UHPLC coupled to an Orbitrap IQ-X Tribrid MS (HESI positive mode) with a Waters Acquity BEH Amide column (150 × 2.1 mm, 1.7 µm) and precolumn. Solvent A was water with 10 mM ammonium formate + 0.1% formic acid; solvent B was acetonitrile + 0.1% formic acid. The gradient ran from 95% to 40% B over 10.5 min, followed by re-equilibration. Source settings included spray voltage 3.5 kV, capillary temperature 300 °C, and vaporizer temperature 275 °C. MS1 parameters: resolution 120,000, AGC target 4e5, scan range 70-800 m/z. Compound identity was confirmed with authentic chemical standards. Data were processed in Skyline (v24.1.1.398) and plotted in GraphPad Prism. Statistical significance was assessed by an unpaired Student’s *t*-test.

#### TBARS assay for MDA estimation

TBARS assay was done following our previously published protocol (cells were plated in 10 cm culture dishes and maintained in complete growth medium for 24 h to allow adherence. They were then exposed to AZD0156 at a final concentration of 0.5 µM. After 72 h of treatment, cells were collected by trypsinization, rinsed with ice-cold PBS, and counted. For each experimental condition, 1 × 10⁷ cells were used to measure MDA content, an indicator of lipid peroxidation, using the TBARS Assay Kit (Cayman Chemical, 10009055) according to the manufacturer’s protocol. Data was plotted using Graphpad Prism software. An unpaired student’s *t*-test was used to determine significance.

### QUANTIFICATION AND STATISTICAL ANALYSIS

#### RNA-seq and analysis

RNA-seq libraries were prepared from 4 separate cell lines (5 replicates each for vehicle or AZD0156 treatment) that comprised the most susceptible SCLC cell lines (H446, H196, DMS114, H720) to AZD0156 treatment. Freshly prepared cell pellets were submitted to GENEWIZ for RNA isolation, library construction, and RNA-seq. Total RNA was extracted using the RNeasy Plus Universal Mini Kit (Qiagen, Hilden, Germany) in accordance with the manufacturer’s instructions. RNA-seq libraries for Illumina sequencing were generated with the NEBNext Ultra II RNA Library Prep Kit (NEB, Ipswich, MA, USA) following the provided protocol. RNA integrity and library quality were assessed using the Agilent TapeStation system (Agilent Technologies, Palo Alto, CA, USA). Library concentrations were determined by both qPCR (KAPA Biosystems, Wilmington, MA, USA) and Qubit 2.0 Fluorometer (Invitrogen, Carlsbad, CA, USA). Sequencing libraries were loaded onto an Illumina HiSeq 4000 (or equivalent platform) after clustering on a single flow cell lane, in accordance with the manufacturer’s guidelines. Sequencing was performed in a 2 × 150 bp paired-end format. Image analysis and base calling were carried out using HiSeq Control Software (HCS), and raw data files (.bcl) were converted to FASTQ format and demultiplexed with Illumina’s bcl2fastq v2.17 software, permitting 1 mismatch in index sequence recognition.

A total of 40 RNA-seq libraries from 4 separate cell lines (5 control replicates and 5 AZD0156 treatment replicates) representing 3 SCLC subtypes were processed using the same pipeline for compatibility. Quality control was performed using FastQC (v0.11.8) (Andrews 2010). Trim Galore! (version 0.6.6) was used to trim the adapter sequences with a quality threshold of 20. The human genome reference used for alignment was GRCh38, with GENCODE release 36 serving as the transcriptome annotation. Samples H446 Control replicate 2 and H446 AZD0156 replicate 4 were removed from the analysis due to poor clustering in the PCA (Principal Component Analysis). The alignment was performed using STAR aligner (v2.7.5b). Salmon (v1.2.1) was implemented to obtain gene-level read counts for all libraries. Remaining samples passed the quality control requirements with > 50% of reads uniquely mapping (> 15 M uniquely mapped reads for each library) using STAR aligner.

#### Differential gene expression and functional analysis

Differential expression analysis was performed using the gene-level read counts and the DESeq2 (v1.28.1) R package. Genes with less than 5 reads in total across all samples were filtered as inactive genes. Genes with an adjusted *P*-value of < 0.05 and absolute log_2_ fold change > 0.5 were considered as differentially expressed. *P*-value was adjusted in DESeq2 using the Benjamin-Hochberg method. The over-representation and GSEA for functional enrichment were both performed using the clusterProfiler R package (v3.16.0). Gene sets used for functional enrichment analysis were obtained from the Molecular Signatures Database (MSigDB) and the KEGG PATHWAY database. Fisher’s exact test was applied to assess the statistical significance of overlap between DEGs (Differentially expressed genes) and genes associated with each term (*P* < 0.05). Terms that are significantly enriched (adjusted *P*-value < 0.05) are indicated in bold and marked with an asterisk.

#### Survival analysis

Real-world OS information was obtained from insurance claims data. OS was calculated from start of treatment with cisplatin until last contact. Patients without contact/claims data for a period of at least 100 days were presumed deceased. Conversely, patients with documented clinical activity within 100 days prior to the latest data update were censored in the analysis. Kaplan-Meier estimates were calculated for molecularly defined patient cohorts. Hazard ratios (HRs) were determined by a Cox proportional hazards model and *P*-values by log-rank test (**Supplementary Fig. S1D**).

Survival analysis was conducted using the IMpower133 clinical dataset, which includes 271 SCLC patients treated with either atezolizumab (Atezo) or placebo. To investigate the prognostic relevance of specific transcriptional programs, we focused on leading-edge genes from 3 hallmark pathways: MYC targets, UPR, and oxidative phosphorylation, as identified in prior enrichment analysis (**Fig. 1G**).

Single-sample GSEA (ssGSEA) was applied to compute pathway activity scores for each patient. These scores were then incorporated into a multivariate Cox proportional hazards model alongside treatment status and SCLC molecular subtype classification to evaluate their association with OS. The analysis was performed using the survival package (v3.2-11) in R.

Statistical significance was assessed for the pathway score terms within the model. For visualization, patients were stratified into quartiles based on their pathway scores, and survival curves were generated using ggsurvplot() from the survminer (v0.4.9) package for the first and fourth quartiles (*n* = 68 per group) to illustrate differences in survival outcomes.

#### CRISPR screen data processing and analysis

CRISPR screen data were processed and analyzed using the MAGeCK (v0.5.9.5) and MAGeCKFlute (v2.9.0) pipelines. Raw paired-end FASTQ files were aligned to a trimmed sgRNA reference library using MAGeCK count, with total normalization applied to account for sequencing depth and sample-specific biases. Non-targeting control sgRNAs were used to estimate background noise and guide normalization.

Quality control metrics, including Gini index (ranging from 0.07-0.15), mapping ratios (> 90% of reads mapped), and dropout rates (∼ 1.4%), were assessed using MAGeCKFlute and visualized with ggplot2 and pheatmap. Heatmaps of raw and normalized counts were generated for all sgRNAs and control sgRNAs to evaluate library complexity and normalization performance. The total normalization method was used for all downstream analysis.

Differential sgRNA enrichment analysis was performed using MAGeCK-MLE, incorporating a design matrix to model experimental conditions. Gene-level beta scores were estimated and adjusted using false discovery rate (FDR) correction. To account for systematic biases, beta scores were further normalized using a cell cycle-based normalization method implemented in MAGeCKFlute, enabling robust cross-condition comparisons. Candidate gene hits were identified using a 9-square plot framework implemented via the SquareView() function. This approach classifies genes based on their beta score enrichment or depletion across conditions. The *x*-axis represented control beta scores at T1 or T2 relative to T0, while the *y*-axis represented treatment beta scores at T1 or T2 relative to T0. Diagonal boundaries reflect a ±1.5 standard deviation cutoff used to identify genes with general positive or negative selection. Vertical and horizontal boundaries represent a ±2 standard deviation cutoff, highlighting genes with treatment-specific effects. Genes located in the top-center and bottom-center quadrants, indicating strong enrichment or depletion under treatment but not control, were prioritized.

#### Metabolomics data processing and analysis

Raw metabolite intensity data were imported and preprocessed in R. Molecules with missing values across all samples were excluded to ensure data integrity. Intensity values were log-transformed using a log10(*x* + 1) transformation to stabilize variance and reduce skewness. The resulting matrix was *Z*-score normalized across samples to center and scale the data for downstream multivariate analysis. Differential abundance analysis of metabolites was performed using the limma (v3.48.0) package in R. For each cell line, log-transformed and *Z*-score normalized metabolite intensities were subset and modeled using linear models. Experimental groups (e.g., control vs. AZD0156-treated) were encoded in a design matrix, and contrasts were specified to estimate treatment effects. Empirical Bayes moderation was applied to improve variance estimation across metabolites. Significance was assessed using moderated *t*-statistics, and *P*-values were adjusted for multiple testing using the FDR method. Metabolites with an adjusted *P*-value < 0.05 and absolute log_2_ fold change > 0.25 were considered significantly differentially abundant. Volcano plots were generated using ggplot2 to visualize differential metabolites, highlighting upregulated and downregulated features.

#### IC_50_ biomarker analysis

To identify gene expression biomarkers associated with sensitivity to ATM inhibition, we analyzed IC_50_ values for AZD0156 across 22 SCLC cell lines. IC_50_ data were curated and matched to transcriptomic profiles from the CellMiner database. Gene expression data were log-transformed and correlated with IC_50_ values using Pearson correlation. Correlation coefficients and *P*-values were computed for each gene, and multiple testing correction was applied using the FDR method. Genes with significant positive or negative correlations were ranked to generate a gene list for pathway-level analysis.

GSEA was performed using the clusterProfiler package and Hallmark gene sets from MSigDB. Pathways with adjusted *P*-values < 0.05 were considered significantly enriched. Core enrichment genes from the top enriched pathways were extracted for further analysis.

To assess pathway-level expression patterns, we computed the mean expression of core enrichment genes across cell lines and visualized these using a heatmap. Cell lines were ordered by increasing IC_50_ values to highlight expression trends associated with drug sensitivity.

#### Heatmaps and boxplots

Heatmaps were generated using the R package pheatmap (v1.0.12) to visualize patterns in drug sensitivity, gene expression, and pathway activity across cell lines. For **Figure 1E**, *Z*-score-normalized IC_50_ values were plotted with cell lines as columns and drugs as rows. Cell lines were ordered by increasing ATM expression, as indicated in the top annotation bar, and Spearman correlation coefficients between ATM expression and drug sensitivity were displayed in the side annotation bar. **Figure 1G** presents the mean expression of core enrichment genes from significant pathways across cell lines, derived from GSEA. **Supplementary Figure 5B** shows beta scores for the top 35 genes associated with susceptibility to AZD0156 treatment at time point T1 in CRISPR-library infected H196 cells, highlighting ACSL1 and VDAC3 as key candidates. **Supplementary Figure 5D** displays RNA-seq expression levels of ferroptosis-related markers, including ACSL1 and VDAC3, in H196 cells treated with an ATMi. All heatmaps were scaled by row to emphasize relative differences and were clustered where appropriate to reveal co-expression patterns.

Boxplots in **Figures 2D, 2L, 3E, 5A-5C,** and **5G** and **Supplementary Figures 3A** and **5E-F** were generated using ggplot2 (v3.4.2). Those in **Figures 1A, 1B, 1D,** and **Supplementary Figure S1C** were generated using JMP® 18.0.1 (NC, USA). Boxplots are defined as the median (center line); Interquartile range (box; 25th–75th percentile), and whiskers extending to the most extreme values within ±1.5×IQR. Boxplots representing pathway signature scores were computed using the ssGSEA method, while boxplots showing individual gene expression reflect normalized expression values.

#### Public ATAC-seq and Cistrome data browser for ChIP-seq

To look at accessibility in the ATF4 promoter, publicly available ATAC-seq data (GEO ID - GSE256345) was downloaded and analyzed for untreated H82 SCLC cell lines. Quality control was performed using FastQC (v0.11.8). Trim Galore! (v0.6.6) was used to trim the adapter sequences with default parameters. For each individual sample, paired-end 150-bp reads were aligned to the human reference genome (GRCh38/GENCODE release 36) using Bowtie2 (v2.2.8) with default parameters and –X 2000. Reads were sorted using SAMtools (v1.11), and mitochondrial and pseudo-chromosomal alignments were removed. Picard (v2.2.4) was used to remove duplicates (Picard Toolkit 2019). Peaks were called using MACS3 (v3.0.0a7) with parameters -nomodel -nolambda -slocal 10000. Coverage tracks (Bigwig files) were generated from filtered BAM files for individual replicates using deepTools (v3.2.1). The bamCoverage tool was run with the following parameters: --normalizeUsing RPKM, binsize 10, and extendReads 200. The coverage tracks were uploaded and visualized on the UCSC Genome Browser.

The Cistrome Data Browser, which hosts publicly available processed ChIP-seq datasets, was leveraged to visualize if MYC was bound to ATF4 in SCLC cell lines. The inputs to the browser were as follows: species – Homo sapiens, assay – ChIP-seq, ontology - small cell lung carcinoma, factor – Myc. Peak calls were derived from pile-up files obtained from the Cistrome database, representing regions of significant signal enrichment.

## Supporting information

Graphical Abstract

## Acknowledgments

This work was supported in part by the Bioinformatics for Next Generation Sequencing (BiNGS) shared resource facility within the Tisch Cancer Institute at the Icahn School of Medicine at Mount Sinai, which is partially supported by NIH grant P30CA196521. This work was also supported in part through the computational resources and staff expertise provided by Scientific Computing at the Icahn School of Medicine at Mount Sinai and supported by the Clinical and Translational Science Awards (CTSA) grant UL1TR004419 from the National Center for Advancing Translational Sciences. Research reported in this paper was supported by the Office of Research Infrastructure of the National Institutes of Health under award number S10OD026880 and S10OD030463. The authors would like to thank Angela Dahlberg, editor in the OSUCCC Division of Medical Oncology, for editing this manuscript.

