## Supplementary Figures for "ATM functions as a rheostat of metabolic stress in small-cell lung cancer"

### Supplementary Figure 1

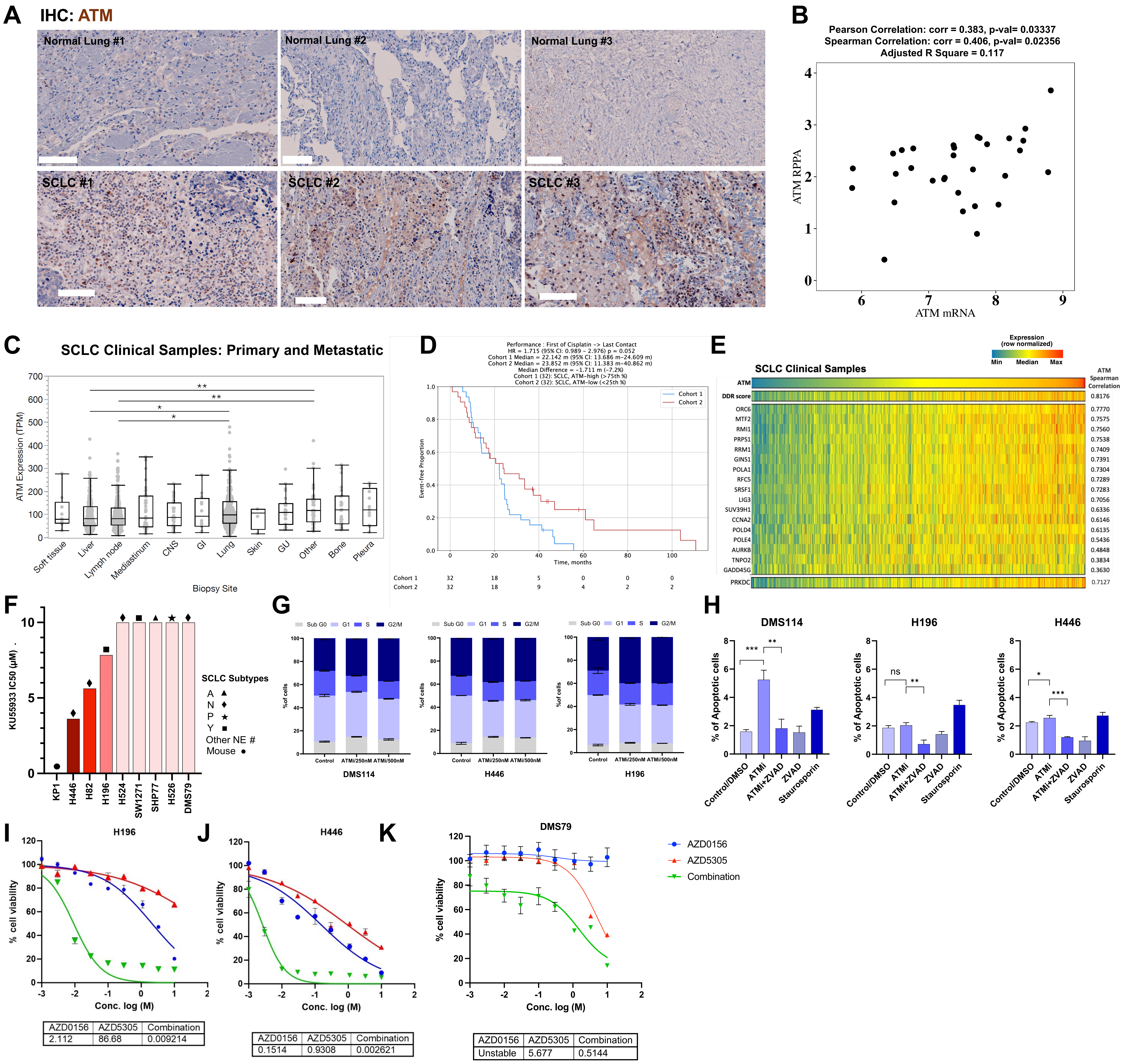

### Supplementary Figure 2

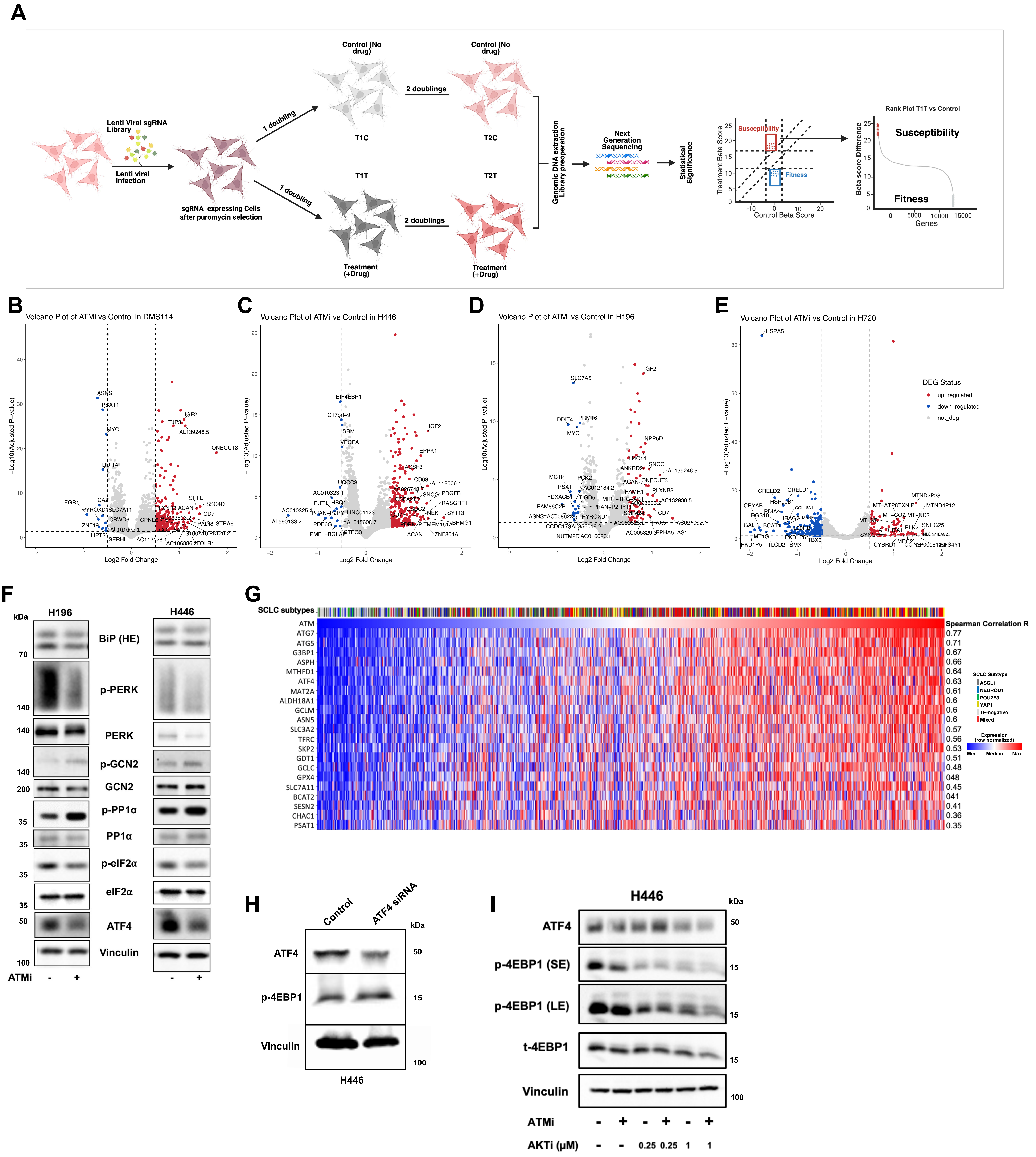

### Supplementary Figure 3

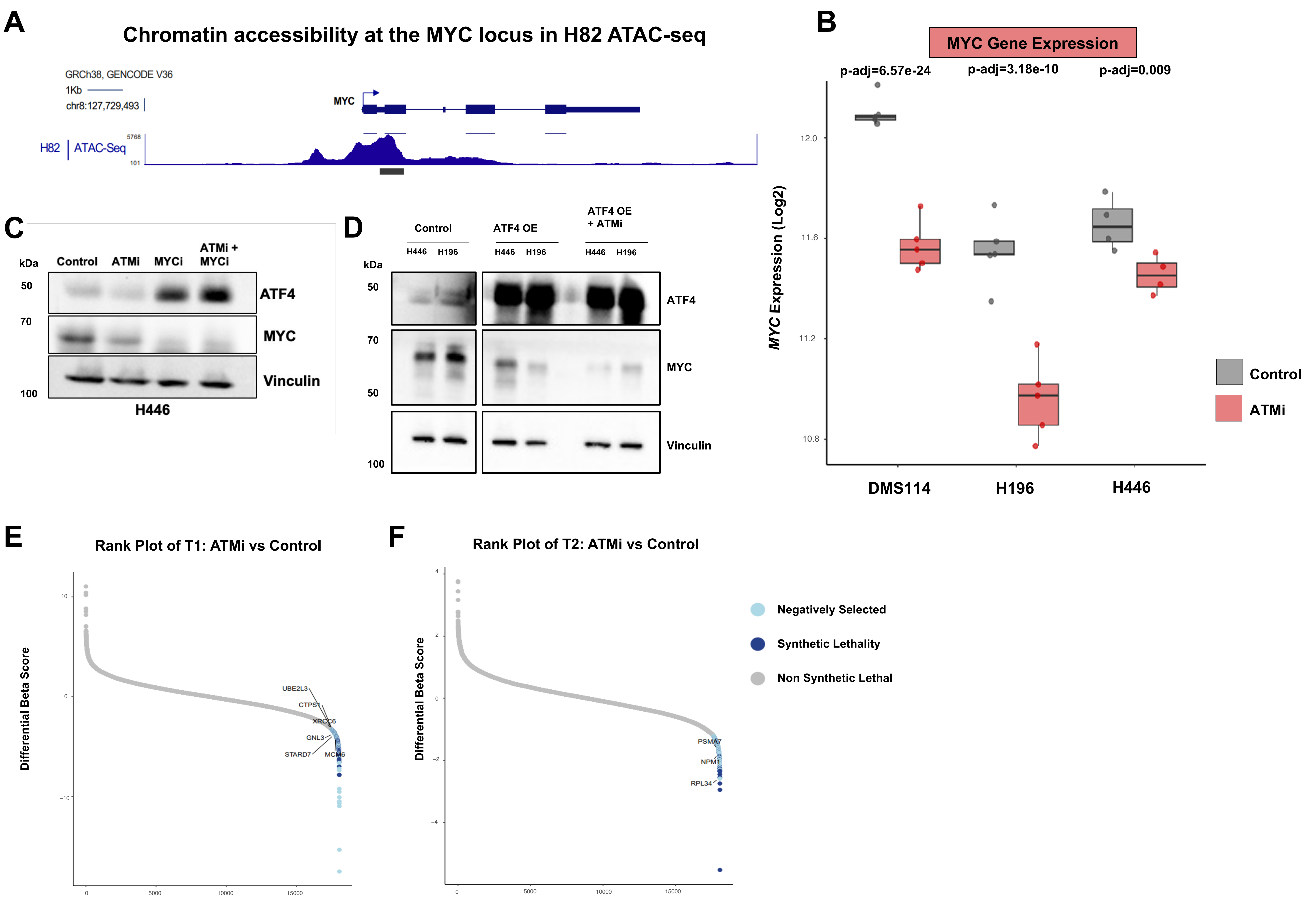

#### Supplementary Figure 4

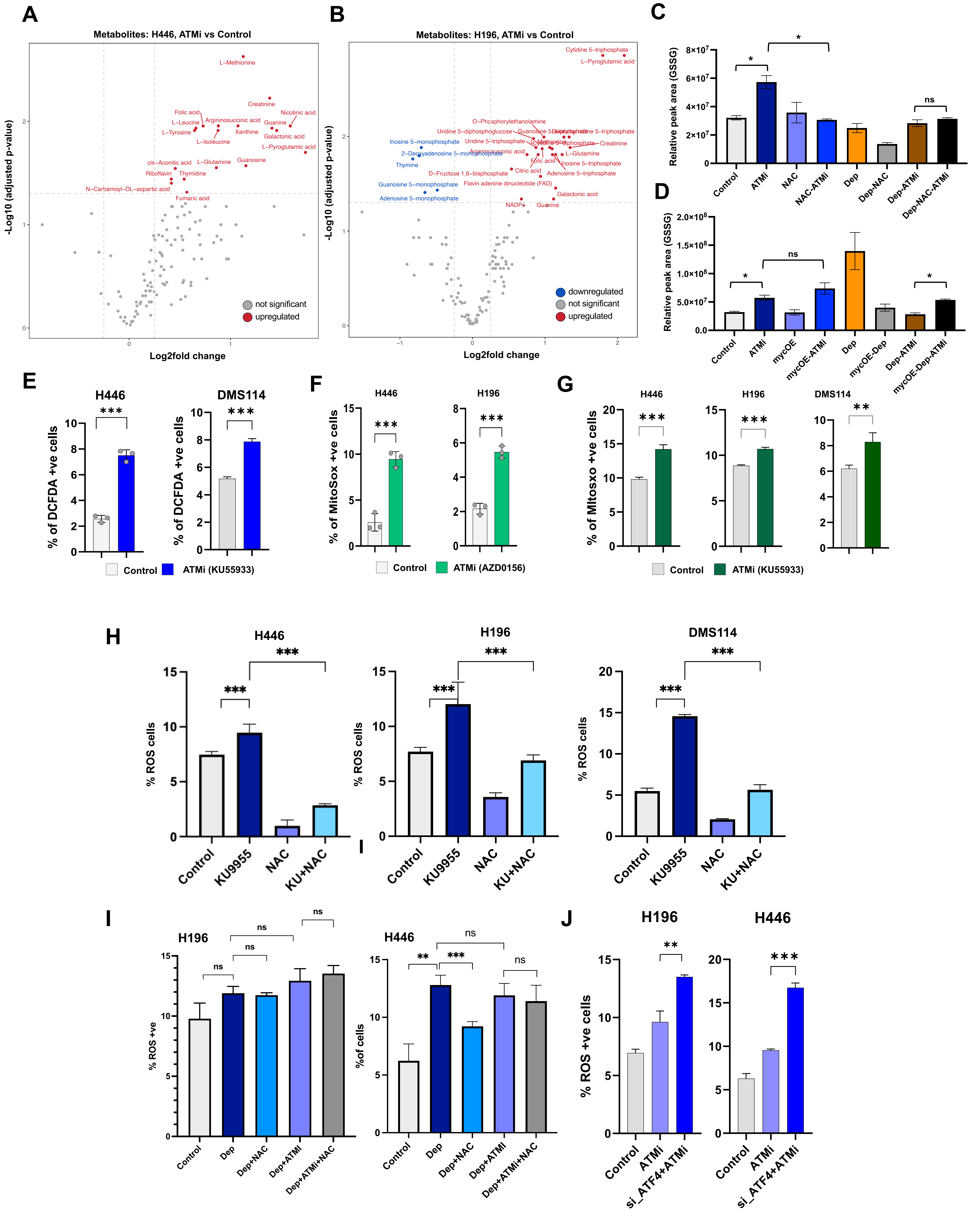

### Supplementary Figure 5

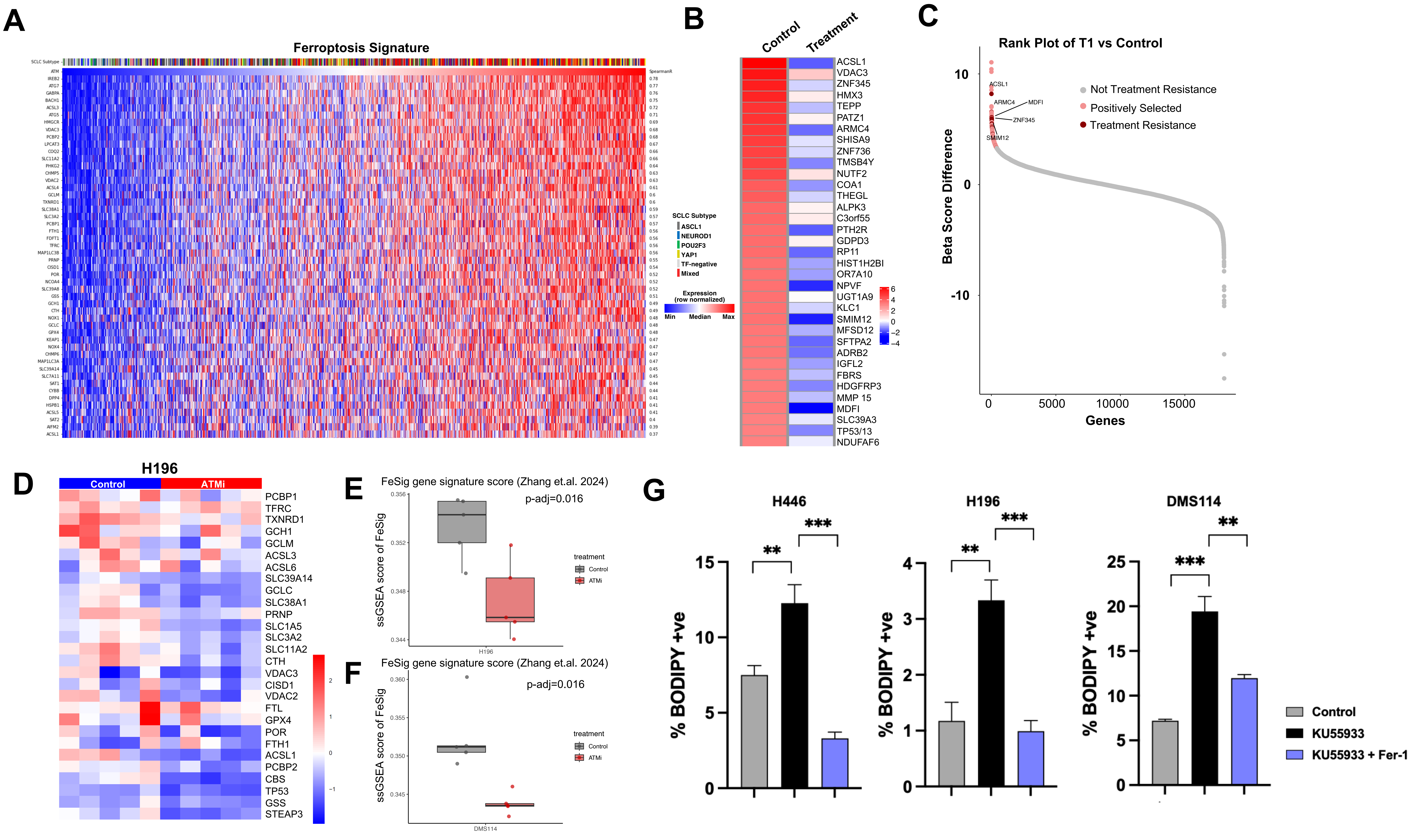

### Supplementary Figure 6

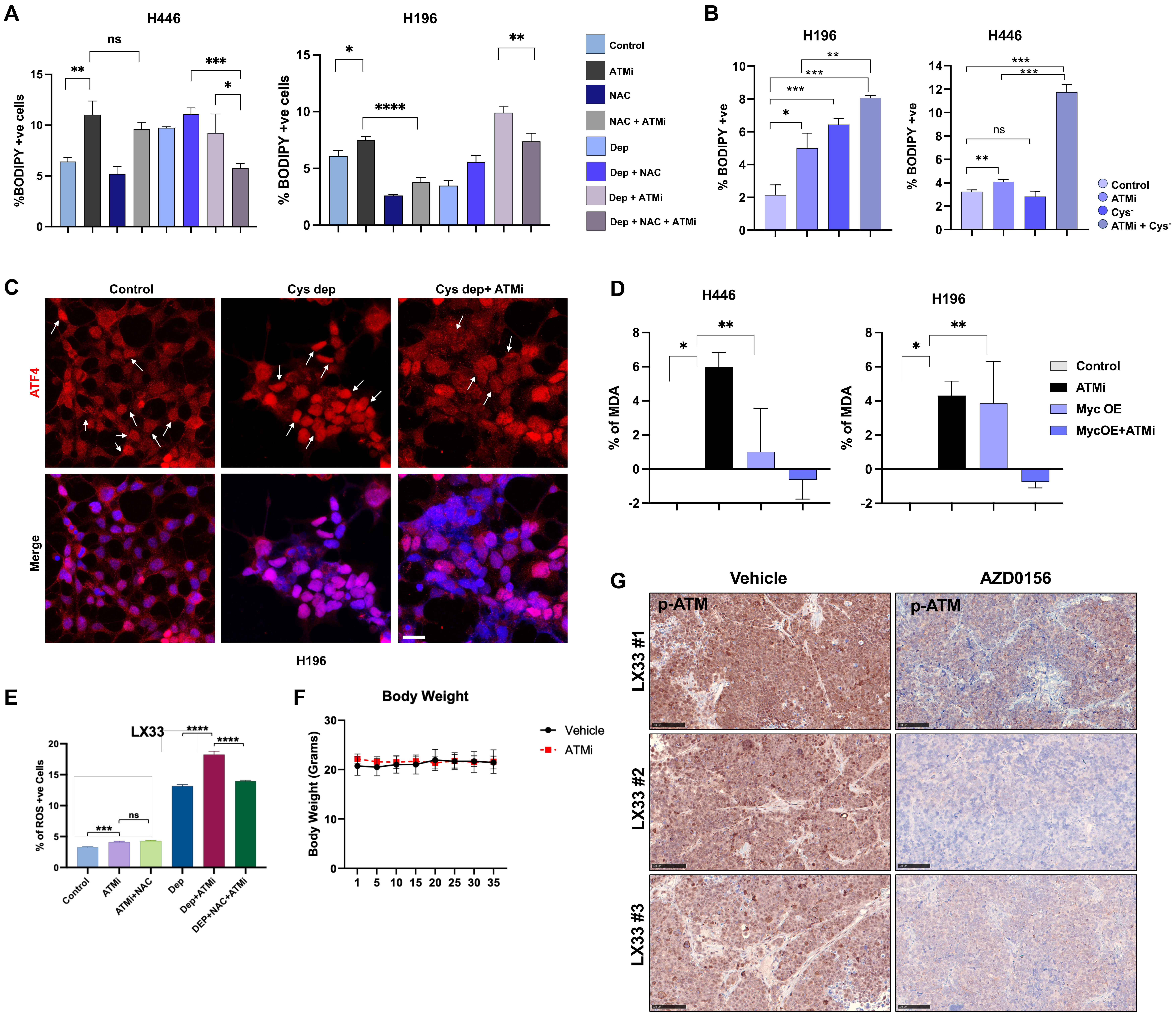
