## Supplementary figures and images for "ATM functions as a rheostat of metabolic stress in small-cell lung cancer"

### Graphical Abstract

#### Graphical Abstract

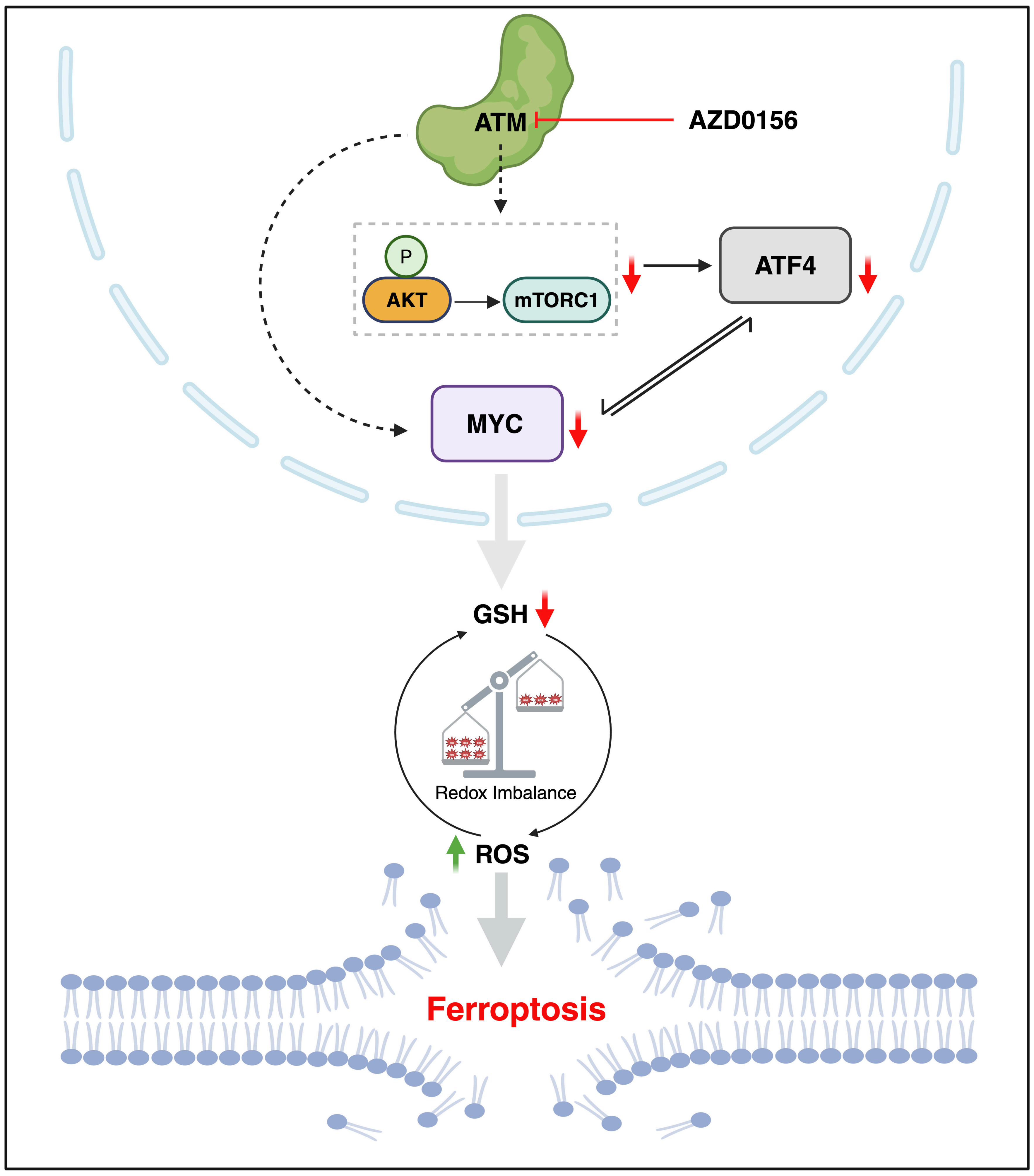
