## Supplementary Figure Legends for "ATM functions as a rheostat of metabolic stress in small-cell lung cancer"

**Supplementary Figure S1**. **ATM in SCLC patient samples and combination strategies,** **Related to Figure 1**

**(A)** IHC of real-world patient tissue with ATM across 3 normal lung and 3 SCLC patient samples. Scalebar = 100 um. **(B)** Correlation between ATM mRNA expression and ATM protein levels across SCLC tumors. mRNA expression (*x*-axis) and protein levels measured by reverse phase protein array (RPPA; *y*-axis) were plotted for matched tumor samples. Each dot represents an individual tumor. A positive trend suggests transcriptional regulation of ATM protein abundance. Correlation coefficient values are reported at the top of the graph. **(C)** SCLC primary and metastatic-sitewise comparison of mRNA expression of ATM in real-world patient data. **(D)** Kaplan-Meier plot showing OS in months, starting from the first administration of cisplatin treatment in patients with SCLC, stratified by median ATM expression levels using a Cox regression model. **(E)** Heatmap of gene expression profiles of DDR genes in real-world patient samples ordered by ATM expression and showing the correlation value and total DDR score. **(F)** IC_50_ values of 9 SCLC cell lines representing major subtypes of SCLC treated with ATM inhibitor KU55933. **(G-H)** Cell cycle **(G)** or apoptosis **(H)** FACS-based assay with annexin-PI of most susceptible SCLC lines H446, H196, and DMS114 +/- ATM inhibition with AZD0156. ZVAD and staurosporin were used for rescue and as a positive control for apoptosis, respectively. **(I-K)** PARPi (AZD5305), AZD0156, or both (combination) in varying concentrations in AZD0156-susceptible cell lines H196 **(I)**, H446 **(J),** and AZD0156-resistant line DMS79 **(K)**. Cell viability was measured using the CellTiter-Glo™ Luminescent Cell Viability Assay. The box underneath each graph represents the IC_50_ value of individual inhibitors or in combination.

**Supplementary Figure S2. Genome-wide CRISPR screen, transcriptomic and phosphoproteomic alterations on ATM inhibition, related to Figure 2**

**(A)** Schematic diagram representing the genome-wide CRISPR screen using the TKVO3 library across 3 doublings. **(B-E)** Volcano plots showing differential gene expression in DMS114 **(B)**, H446 **(C)**, H196 **(D),** and H720 **(E).** SCLC cells treated with ATM inhibitor AZD0156 compared to DMSO control. Log₂ fold change is plotted on the *x*-axis and -log₁₀ (adjusted *P*-value) on the *y*-axis. Red and blue dots represent significantly upregulated and downregulated genes, respectively (adjusted *P*-value < 0.05 and |log₂FC| > 0.5). Selected genes of interest are labeled. **(F)** Western blot panel of H446 and H196 +/- ATMi (AZD0156) probed with UPR markers BiP, p-PERK, PERK, p-GCN2, GCN2, p-PP1α, PP1α, p-eIF2⍺, eIF2⍺, ATF4, and vinculin. **(G)** Heatmap showing expression levels of ATF4 targets: stress-response and redox-regulating genes across 944 real-world SCLC patient tumor samples. Rows represent genes and columns represent individual tumor samples. Expression values are *Z*-score-normalized by gene (row), with red indicating higher and blue indicating lower relative expression. Samples are ordered by increasing ATM expression (top color bar). The uppermost color bar denotes clinical specimen subtype. **(H)** Western blot of H446 treated with or without ATF4 siRNA and probed for p-4EBP1, ATF4, and vinculin. **(I)** Western blot analysis of H446 cells treated +/- ATMi (1 uM) and +/- AKTi and probed for ATF4, p-4EBP1, 4EBP1, and vinculin.

**Supplementary Figure S3. MYC and ATF4 coregulation by ATM in SCLC cell lines, related to Figure 3**

**(A)** ATAC-seq chromatin accessibility track for the MYC gene locus in H82 SCLC cells. Gene orientation and structure are shown (UCSC Genes track), with the MYC gene transcribed left-to-right on chromosome 22 (chr22:39,517,624–39,528,368). Blue bars denote annotated exons. **(B)** Boxplot showing MYC gene expression in SCLC cells treated with ATM inhibitor AZD0156. (red) compared to DMSO controls (gray) across three SCLC cell lines - DMS114, H196 and H446. P-values were calculated using a two sample t-test and were adjusted using the BH method (P-adj: DMS114 – 6.57e-24, H196 – 3.18e-10 and H446 - 0.00955)**(C)** Western blot analysis of H446 treated with MYCi +/- ATMi and probed for ATF4, MYC, and vinculin (loading control). **(D)** Western blot of H446 and H196 cells with/without transient ATF4 overexpression +/- ATMi probed for ATF4 and MYC. Vinculin was probed as a loading control. **(E-F)** Beta score-essentiality sigmoid plot from CRISPR analysis at T1 **(E)** and T2 **(F)**. Genes contributing to *MYC* targets are enclosed in boxes.

**Supplementary Figure S4. Metabolic regulation of SCLC by ATM-MYC signaling, related to Figure 4**

**(A-B)** Volcano plots depicting differentially abundant metabolites in H446 **(A)** or H196 **(B)** treated with ATMi (AZD0156, 72 h) versus control. The *x*-axis represents log₂ fold change, and the *y*-axis represents -log₁₀ (adjusted *P*-value). Red and blue dots indicate significantly upregulated and downregulated metabolites, respectively (*P* < 0.05, |log₂FC| > 0.25), and gray dots represent metabolites that were not significantly changed. **(C-D)** Relative intracellular GSSG levels quantified by targeted LC-MS +/- ATMi +/- NAC **(C)** or +/- ATMi and/or stably overexpressing MYC **(D)**, cultured in complete or depleted media. **(E)** Relative intracellular ROS measured by DCFDA positive staining in FACS +/- ATMi (KU55933) in H446 and DMS114. **(F-H)** Relative mitochondrial ROS measured by MitoSOX positive staining in FACS +/- ATMi (AZD0156) **(F)**, MitoSOX positive staining in FACS +/- ATMi (KU55933), and intracellular ROS measured by DCFDA positive staining in FACS +/- ATMi (KU55933) +/- NAC **(H)** in SCLC models and **(I)** cultured in depleted media. **(J)** Barplot representing relative intracellular ROS by DCFDA staining +/- ATMi (AZD0156) with transient knockdown of ATF4 by siRNA in H446 and H196. Data represent mean ± SEM of *n* = 3 biological replicates. Statistical significance was determined by 1-way student’s *t*-test (**P* < 0.05; ** *P* < 0.01; *** *P* < 0.001, **** *P* < 0.0001).

**Supplementary Figure S5. ATM inhibition causes ferroptosis, related to Figure 5.**

**(A)** Heatmap showing expression levels of ferroptosis-related genes across 944 real-world SCLC patient tumor samples. Rows represent genes, and columns represent individual tumor samples. Expression values are *Z*-score-normalized by gene (row), with red indicating higher and blue indicating lower relative expression. Samples are ordered by increasing ATM expression (top color bar). The uppermost color bar denotes clinical specimen subtype. **(B)** Top 35 genes associated with susceptibility at T1 on AZD0156 treatment of CRISPR-library infected H196. Note that ACSL1 and VDAC3 are at the top of the list. **(C)** Beta score-essentiality sigmoid plot from CRISPR analysis at T1. **(D)** Heatmap representing gene expression (RNA-seq) of ferroptosis markers including VDAC3 and ACSL1 in H196 treated with ATMi (AZD0156). **(E-F)** Boxplots depicting prognostic signature FeSig score coined by Zhang et Al., 2024 in H196 **(E)** and DMS114 **(F). (G)** Barplot representing relative lipid peroxidation at 24 h measured by BODIPY/C11 positive staining in FACS +/- ATMi (KU55933) in SCLC models with or without ferrostatin-1 (ferroptosis inhibitor). Statistical significance was determined by 1-way student’s *t*-test (**P* < 0.05; ** *P* < 0.01; *** *P* < 0.001, **** *P* < 0.0001).

**Supplementary Figure S6. Lipid peroxidation in-vitro and in-vivo is regulated by ATM expression, related to Figure 5**

**(A)** Lipid peroxidation measurement by BODIPY/C11 in H446 and H196 treated with ATMi (AZD0156) with or without NAC supplementation in complete or minimal/depleted (dep) media. Data represent mean ± SEM of *n* = 3 biological replicates. **(B)** Lipid peroxidation measurement by BODIPY/C11 in H446 and H196 treated with ATMi (AZD0156) for 24 h with or without cysteine depletion for 12 h. **(C)** Confocal images of H196 +/- ATMi showing immunofluorescence for ATF4 in cysteine-depleted growth media conditions. DAPI was stained to visualize nuclei. White arrows point to nuclear accumulation of ATF4. Scalebar: 25 µM. **(D)** Barplot representing relative malonedialdehyde (MDA) content in whole-cell lysates of H446 and H196 treated with ATMi (AZD0156) and/or stably overexpressing MYC. **(E)** Barplot depicting relative intracellular ROS by %DCFDA staining by FACS in PDX-derived cells LX33 in complete and depleted media +/- ATMi, with or without NAC supplementation, in vitro. Statistical significance was determined by 1-way student’s *t*-test (**P* < 0.05; ***P* < 0.01; *** *P* < 0.001, **** *P* < 0.0001). **(F)** Graph depicts body weight of mice over time treated with vehicle or ATMi (AZD0156), with tumor generated subcutaneously from LX33. Data is represented as mean ± SEM (*n* = 6 per group). **(G)** IHC images of LX33 tumors +/- AZD0156 probed for expression of p-ATM, the active form of ATM. 50 mg/kg AZD0156 was the dosage 5/7 days, given via intratumoral injection.
